# Spinal Cord Injury Imprints a Glucocorticoid-Driven Failure State in Hematopoietic Stem Cells

**DOI:** 10.1101/2025.10.05.680535

**Authors:** Kyleigh A. Rodgers, Elizabeth A.R. Garfinkle, Kristina A. Kigerl, Elham Asghari Adib, Katherine A. Mifflin, Jodie Hall, Rohan Kulkarni, Cankun Wang, Chinmayee Goda, Ana C Rodrigues Dias, Zhen Guan, Malith Karunasiri, Qin Ma, Katherine E. Miller, Adrienne M. Dorrance, Phillip G. Popovich

## Abstract

Spinal cord injury (SCI) triggers systemic pathology beyond the nervous system, including bone marrow failure that can worsen infection risk, anemia, and motor recovery. Here we identify a neuroendocrine mechanism that rapidly imprints long-lasting dysfunction in hematopoietic stem cells (HSCs), which sustain lifelong production of immune cells, red blood cells, and platelets. Rather than mounting a canonical stress-hematopoietic response, SCI HSCs enter a broadly repressed state marked by chromatin closure and suppression of programs required for cell-cycle entry, genome maintenance, and redox defense. This maladaptive state leads to persistent DNA damage, impaired oxidative stress resolution, pancytopenia, and loss of long-term HSC regenerative capacity. Mechanistically, SCI-induced glucocorticoid surges drive glucocorticoid receptor-dependent repression of DNA repair genes, including *Lig1* and *Fen1*. Acute glucocorticoid receptor blockade after SCI restores durable hematopoiesis, revealing an early therapeutic window to preserve hematopoietic integrity after neurotrauma.

**Graphical abstract:** 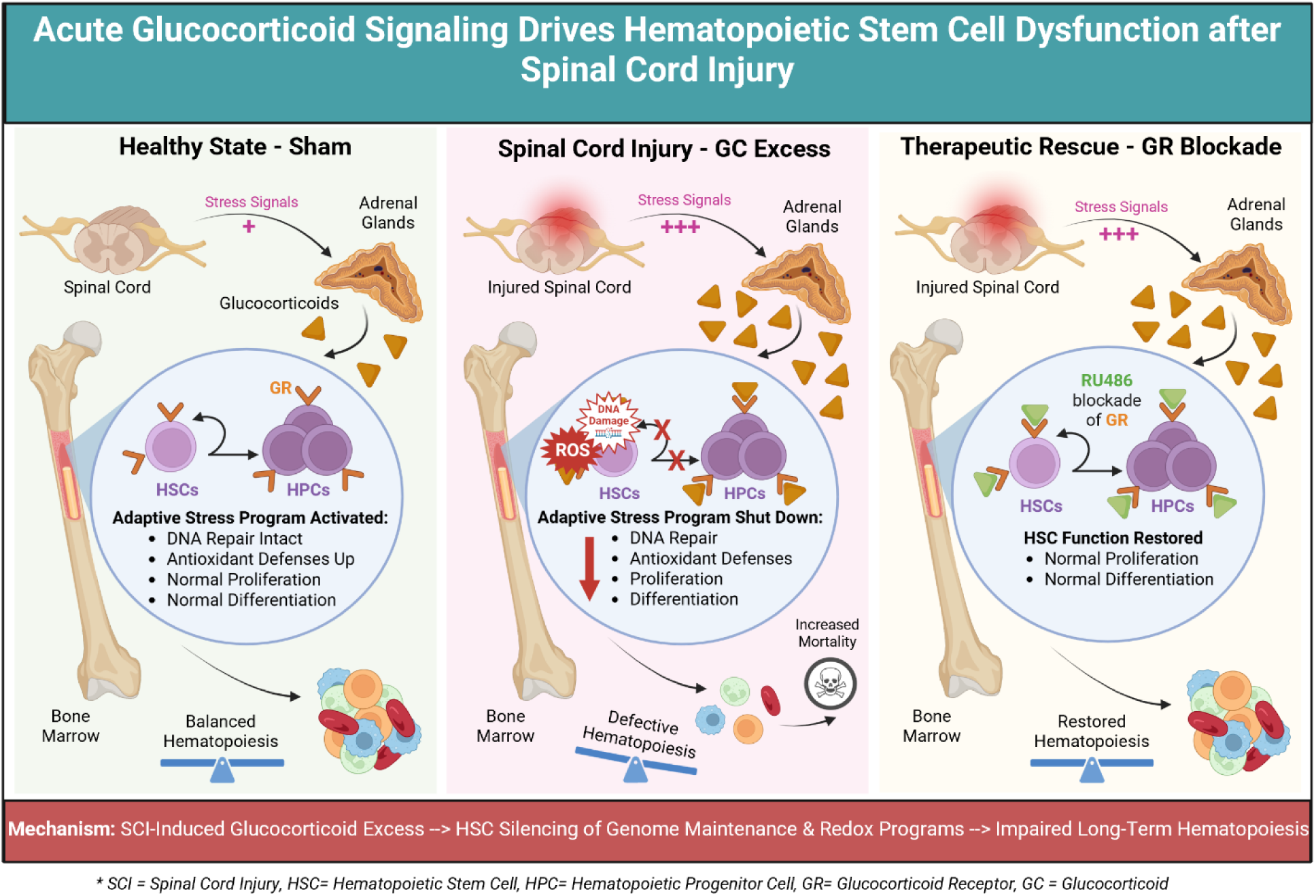

**Key Findings:**

- Acute SCI prevents induction of normal stress-induced repair programs in HSCs.
- SCI HSPCs exhibit increased chromatin compaction.
- SCI compromises HSC genome-maintenance and redox capabilities.
- SCI HSCs have defects in normal cell cycle regulation, are hypersensitive to genotoxic stress, and are deficient in long-term reconstitution.
- Early glucocorticoid receptor (GR) blockade restores long-term hematopoiesis, supporting a GR-dependent but reversible repression mechanism.

## Introduction

Hematopoiesis is a continuous, tightly regulated process generating mature blood cells, primarily within the bone marrow. A hierarchical system begins with hematopoietic stem cells (HSCs), which self-renew and differentiate into hematopoietic progenitor cells (HPCs) ^1,2^. HSCs sustain lifelong hematopoiesis through self-renewal and multilineage differentiation ^3,4^. In contrast, HPCs, derived from HSCs, possess limited self-renewal but robust proliferative capacity, enabling rapid expansion and differentiation into mature blood cells ^5,6^. This division of function allows HSCs to maintain long-term hematopoietic output, while HPCs support short-term production, especially during infection, injury, or inflammation ^7,8^.

HSCs and HPCs, collectively known as HSPCs, regulate blood cell production in response to the body’s needs by adapting to physiological and pathological stress ^9,10^. During disruptions to homeostasis, HSPCs exit quiescence and increase proliferation, which heightens cellular metabolism and generates reactive oxygen species (ROS) and byproducts that may cause DNA damage ^11–13^. Normally, these adaptive responses preserve hematopoietic integrity and restore balance ^14^. However, chronic or severe stress results in excessive ROS-induced DNA damage, triggering cell-cycle arrest and apoptosis, impairing HSC self-renewal and differentiation, and reducing hematopoietic output ^14,15^. This manifests clinically as bone marrow failure, cytopenia, and increased vulnerability to infection, i.e., key features of hematopoietic collapse ^16^.

Central nervous system injuries, including spinal cord injury (SCI), stroke, and traumatic brain injury, disrupt hematopoiesis, resulting in significant hematologic and immune complications ^17–19^. SCI specifically induces chronic immune dysfunction, anemia, increased susceptibility to infection, and impaired wound healing ^20–24^. Persistent inflammation and defective platelet function further exacerbate vascular dysfunction and hinder recovery ^25–27^. The mechanisms by which SCI alters hematopoiesis remain incompletely defined. Although SCI rapidly impairs HSPC function, the cellular basis and potential for therapeutic intervention are not established ^19^. This study aims to address these critical gaps.

New data indicate that as early as one day post-injury, before overt HSPC dysfunction, HSCs from SCI mice display significantly reduced gene expression, increased DNA damage, and disrupted ROS regulation. These intrinsic defects arrest SCI HSCs in the cell cycle. Upon serial transplantation into irradiated healthy recipients, SCI HSCs fail to reconstitute hematopoiesis, confirming cell-intrinsic defects caused by SCI persist indefinitely. Glucocorticoid receptor (GR) signaling can restrain HSC activation by limiting programs required for genome maintenance. SCI triggers a rapid systemic glucocorticoid surge, suggesting a direct endocrine route to the early HSC defects we observe after injury. Accordingly, GR antagonism within 6 h prevents HSC impairment.

These findings demonstrate that SCI rapidly and profoundly compromises HSC function. Targeting HSC dysfunction may represent a novel therapeutic strategy to restore hematopoiesis and improve recovery and long-term outcomes following SCI or other neurological disorders.

## Results

### Spinal cord injury impairs bone marrow hematopoietic stem cell function and lineage commitment

SCI quickly alters circulating leukocytes and is an early post-injury sign of immune dysfunction ^19,28–30^. To assess if these changes start with bone marrow disruption, we used single-cell RNA sequencing (scRNA-seq) to analyze HSPC transcriptional profiles in c-Kit^+^ bone marrow cells from SCI mice at 1 day post-injury (dpi), compared with time-matched sham-operated and naïve controls.

Sham mice were anesthetized and received a laminectomy (removal of bone overlying the spinal segment with associated muscle and skin dissection) but their spinal cords were not injured. This control group helped differentiate cellular responses due to SCI from those caused by surgery. Naïve mice were only anesthetized, with no surgical procedures performed.

Unsupervised clustering of scRNA-seq data from 16,431 bone marrow cells identified 18 distinct HSPC clusters (Fig. 1a). Cell type annotation was based on canonical marker gene expression including established lineage markers (Fig. S1a-c) ^31–34^.

**Fig 1.**
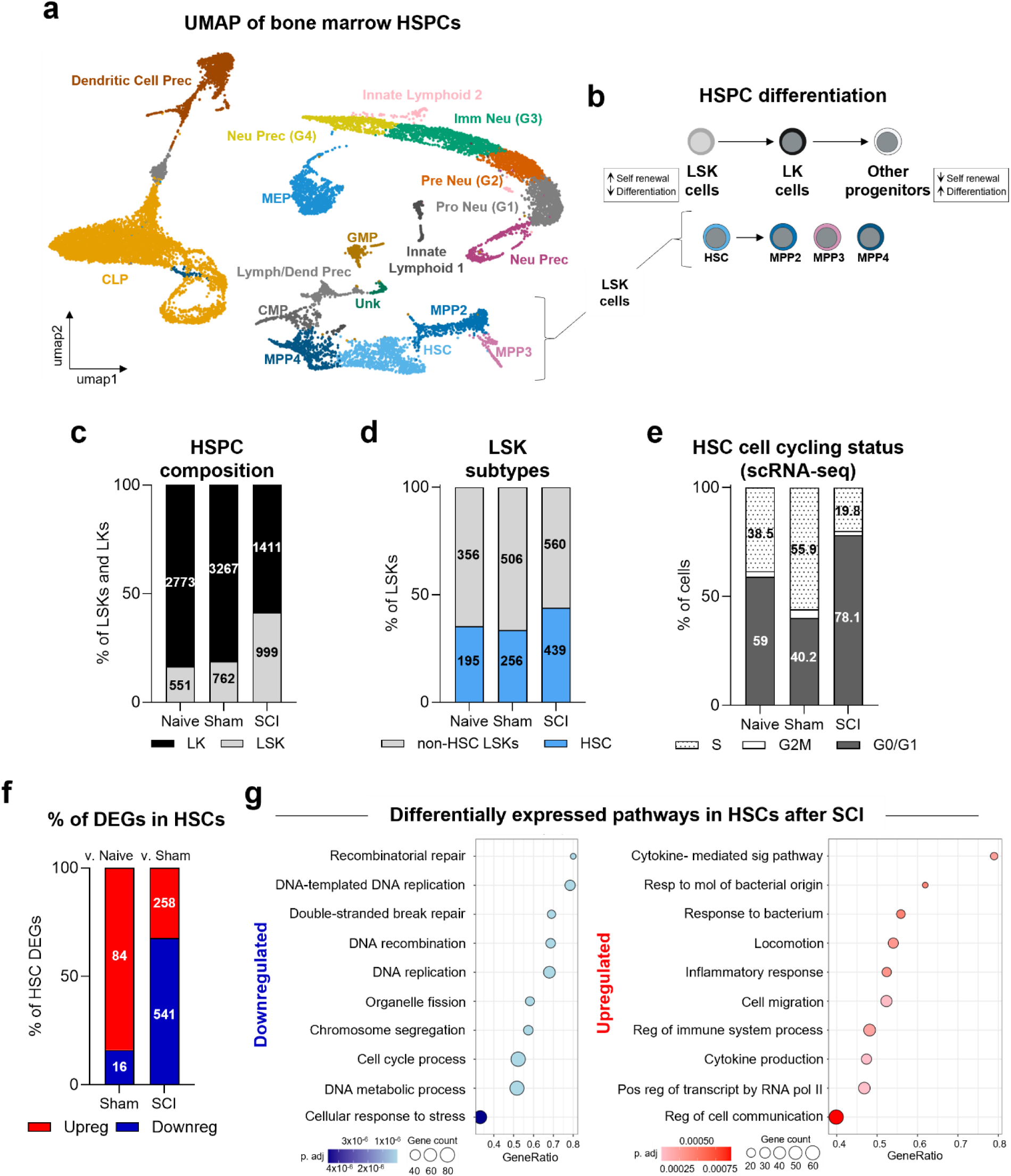
scRNA-seq of bone marrow reveals impaired stress response in HSCs after SCI. Bone marrow HSPCs from naïve, sham, and spinal cord injury (SCI) mice were analyzed by scRNA-seq at 1 day post-injury (dpi) to assess early transcriptional responses to SCI and surgical stress. **(a)** UMAP projection of magnetically selected c-Kit^+^ bone marrow cells from naïve, sham, and SCI mice at 1 dpi. Cells were obtained from one pooled sample per group, with each pool comprising cells from n= 5 mice. LSK cells in brackets. **(b)** Schematic of HSPC functional classes: LSK cells (Lineage⁻ Sca-1⁺ c-Kit⁺) and LK cells (Lineage⁻ Sca-1⁻ c-Kit⁺). LSK cells differentiate into LK cells, which then derive downstream precursors (other). **(c)** scRNA-seq clusters were classified as LSK cells (HSCs and MPPs2-4) and LK cells (GMPs, CLPs, MEPs, and CMPs) and represented as a % of total LSK and LK cells, showing relative population distributions across groups. Numbers represent the cell counts used to calculate percentages. **(d)** % LSK subtypes were classified as HSCs or non-HSCs (i.e., MPPs2-4). Cell numbers shown were used to calculate percentages. **(e)** Cell cycle analysis of HSCs at 1 dpi (dotted: S; white: G2M; grey: G0/G1), showing differences in cell cycle distribution across groups. Numbers represent the % of cells in each phase. **(f)** % of downregulated (blue) and upregulated (red) DEGs in HSCs from sham vs. naïve (left) and SCI vs. sham (right) mice, indicating transcriptional activation following sham surgery and suppression after SCI. Numbers represent the DEGs used to calculate percentages. **(g)** Pathway enrichment visualization for downregulated (blue) and upregulated (red) processes. The GeneRatio reflects the proportion of genes mapping to each pathway and highlights reduced enrichment of stress-responsive and adaptive pathways after SCI. Overall, these data show transcriptional suppression in HSCs after SCI.

To establish a baseline for evaluating the effects of SCI, we first analyzed the effects of sham surgery, a stressor that activates hematopoiesis ^19^. Sham surgery induced a normal canonical stress response compared to naïve controls, marked by expansion of both LSK and LK cells (Fig. 1b,c), likely reflecting increased HSPC proliferation and differentiation ^9,35^. However, the proportional distribution of HSCs and MPPs2-4 within the LSK compartment remained unchanged in sham mice compared to naïve mice (Fig. 1d), suggesting that surgical stress does not significantly alter early lineage composition. Conversely, SCI increased the frequency and number of LSK cells, while reducing LK cells (Fig. 1c). Consistent with prior findings, HSCs were enriched after SCI (Fig. 1d) ^19^. This enrichment suggests altered HSC activation, potentially reflecting impaired differentiation from HSCs to MPPs and changes in HSC proliferation. This increase in LSK cells after SCI therefore likely reflects accumulation rather than increased proliferation at this timepoint. Indeed, SCI HSCs showed reduced expression of genes associated with cycling cells. While over 50% of sham HSCs were in S phase-indicating increased stress-induced cycling, only ∼20% of SCI HSCs were in S phase, with the majority (∼80%) in G0/G1 phase (Fig. 1e).

Changes in cell cycle score coincided with marked transcriptional differences. In sham HSCs, 84% (84/100) of differentially expressed genes (DEGs) were upregulated (Fig. 1f), and known to have roles in metabolic, survival, and proliferative functions. Pathway enrichment analysis was not statistically significant (p-adj > 0.05), indicating that gene expression changes were dispersed across multiple pathways or limited to specific genes within these pathways ^36^. In contrast, SCI HSCs exhibited widespread transcriptional suppression, with approximately 68% (541/799) of DEGs downregulated compared to sham HSCs (Fig. 1f). These downregulated genes regulate DNA strand break repair, chromosomal organization, cell cycle progression, and DNA replication (Fig. 1g) ^13,16^. Similar patterns and composition of cell cycle suppression and broader transcriptional suppression were observed in non-HSC LSK cells (i.e., MPPs) (Fig. S1d,e). Genes that were upregulated by SCI were significantly enriched in immune activation, transcriptional regulation, cellular trafficking, and membrane dynamics (Fig. 1g) ^37,38^.

### SCI disrupts bone marrow HSPC proliferation and differentiation

Since scRNA-seq identified changes in genes regulating HSPC proliferation and differentiation, we quantified HSPC cell cycle status using flow cytometry and Ki67 and DNA content staining (Fig. 2a,b). SCI significantly reduced the percentage of proliferating (Ki67^+^) HSPCs compared to sham controls. The proportion of HSCs in G1 phase increased relative to S-G2-M, indicating a cell cycle arrest at G1 (Fig. 2c-g). Independent validation with BrdU incorporation and immunophenotyping supported these results (Fig. S2). The MPP-to-HSC ratio, a measure of differentiation dynamics, decreased in SCI mice versus naïve and sham groups (1.28 SCI vs. 1.83 naïve and 1.97 sham), primarily due to increased HSCs, demonstrating impaired transition from HSCs to MPPs (Fig. 2h) ^39^. Sham-induced HSPC proliferation correlated with enhanced proliferative capacity assessed by an increase in colony forming unit-assays (CFU) compared to naïve (Fig. 2i). In contrast, there was little evidence of increases in HSPC expansion within SCI bone marrow (Fig. 2i).

**Fig 2.**
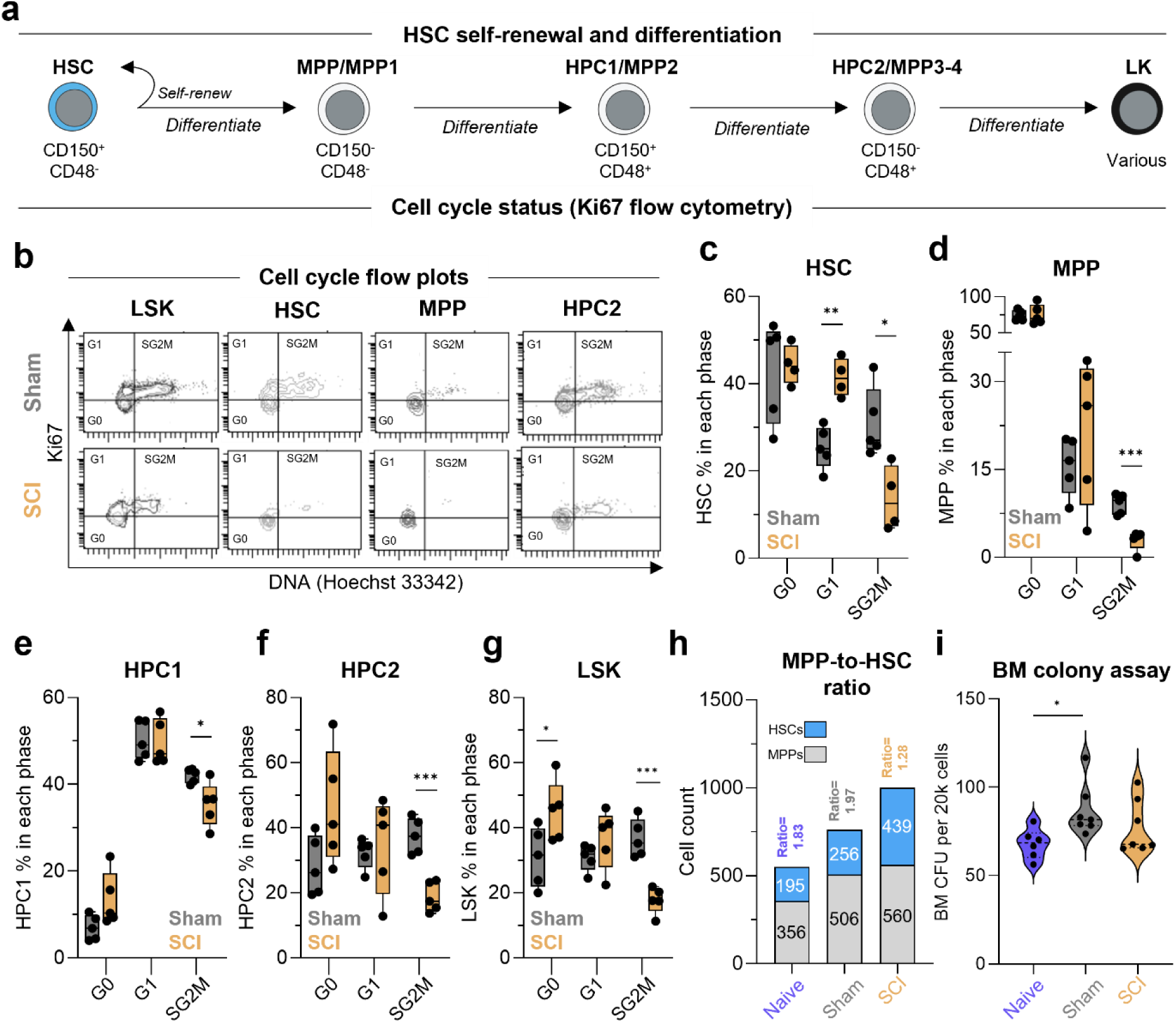
SCI prevents stress-induced HSC proliferation. **(a)** Schematic of hematopoiesis demonstrating that HSCs self-renew or differentiate into downstream progenitors distinguished by flow cytometry cell surface markers shown. **(b)** Representative select flow plots from data in (c-g). Gated on the cell populations labeled. **(c-g)** % of Ki67^+^ HSPC subtypes and nuclear DNA stain (Hoechst 33342) by flow cytometry, demonstrating reduced proliferation with G0 and G1 accumulation in SCI HSPCs compared to sham. **(h)** The y-axis shows the numbers of HSC and non-HSC LSK cells (MPPs2-4) in each sample. The MPP-to-HSC ratio above each bar reflects the relative abundance of downstream progenitors compared to HSCs. The values within each bar indicate the number of cells analyzed in each subtype, illustrating how each population contributes to the overall ratio. **(i)** Colony Forming Unit (CFU) data at 1 dpi in whole bone marrow (WBM) plated in MethoCult M3434. Total colonies were enumerated per 20k WBM cells. Overall, these data show that SCI impairs HSPC proliferation, consistent with suppression of early HSPC activation. **Experimental repetitions**: Data in (c-g, i) are from one experiment that was independently repeated at least twice with similar results. Data in (h) are derived from the same mice as the scRNA-seq dataset (Fig. 1). **Sample sizes:** Mice per group were as follows: n= 4-5 (c-g), n= 6-7 (i), and a single sample per group containing pooled bone marrow from n= 5 mice per group (h). **Statistical analysis:** Box-and-whisker plots (c-g) show the interquartile range (25-75%), with the median indicated by the horizontal line. Whiskers span the full data range. (i) Violin plot shows the median (dotted line). Individual data points by mouse are overlaid. Welch’s t-tests were used for pairwise comparisons between SCI and sham groups within each cell-cycle phase in (c-g). Kruskal-Wallis tests were used for multi-group nonparametric CFU comparisons in (i). *p<0.05, **p<0.01, ***p<0.001. **Exclusion:** One SCI mouse was excluded in (c) due to a staining artifact in the end gate.

Together, data in Figs. 1&2 indicate that SCI rapidly induces defects in the primitive bone marrow HSPCs, distinct from the normal stress responses associated with surgery or anesthesia. Prior work has reported increased HSPC proliferation at later timepoints after SCI (e.g., 3 dpi) ^19^. Current data, obtained at 1 dpi, indicate that SCI initially suppresses normal stress-induced HSC activation and differentiation. Thus, the increased proliferation reported at later time points (e.g., 3 dpi) likely represents a delayed, dysregulated response.

### SCI impairs long-term hematopoietic stem cell function

To determine whether the early transcriptional repression and cell-cycle arrest induced by SCI translate into durable, cell-intrinsic functional deficits, we performed serial *in vivo* competitive repopulation assays, the gold standard for evaluating HSC regenerative capacity (Fig. 3a) ^40^. One day after sham surgery or SCI, donor bone marrow cells (CD45.2^+^) were mixed in equal ratios with sex- and age-matched congenic naïve competitor bone marrow cells (CD45.1^+^) and transplanted into lethally irradiated recipient mice (CD45.1^+^). While short-term hematopoiesis remained intact (Fig. 3b), SCI donor cells exhibited significantly reduced HSC chimerism in bone marrow at 8 months, indicating impaired HSC long-term repopulating capacity (Fig. 3c, S3a). In contrast, progenitor chimerism (MPP, HPC1, and HPC2) remained stable or increased modestly, suggesting that progenitor cells, not HSCs, drive initial short-term engraftment (Fig. 3b,c).

**Fig 3.**
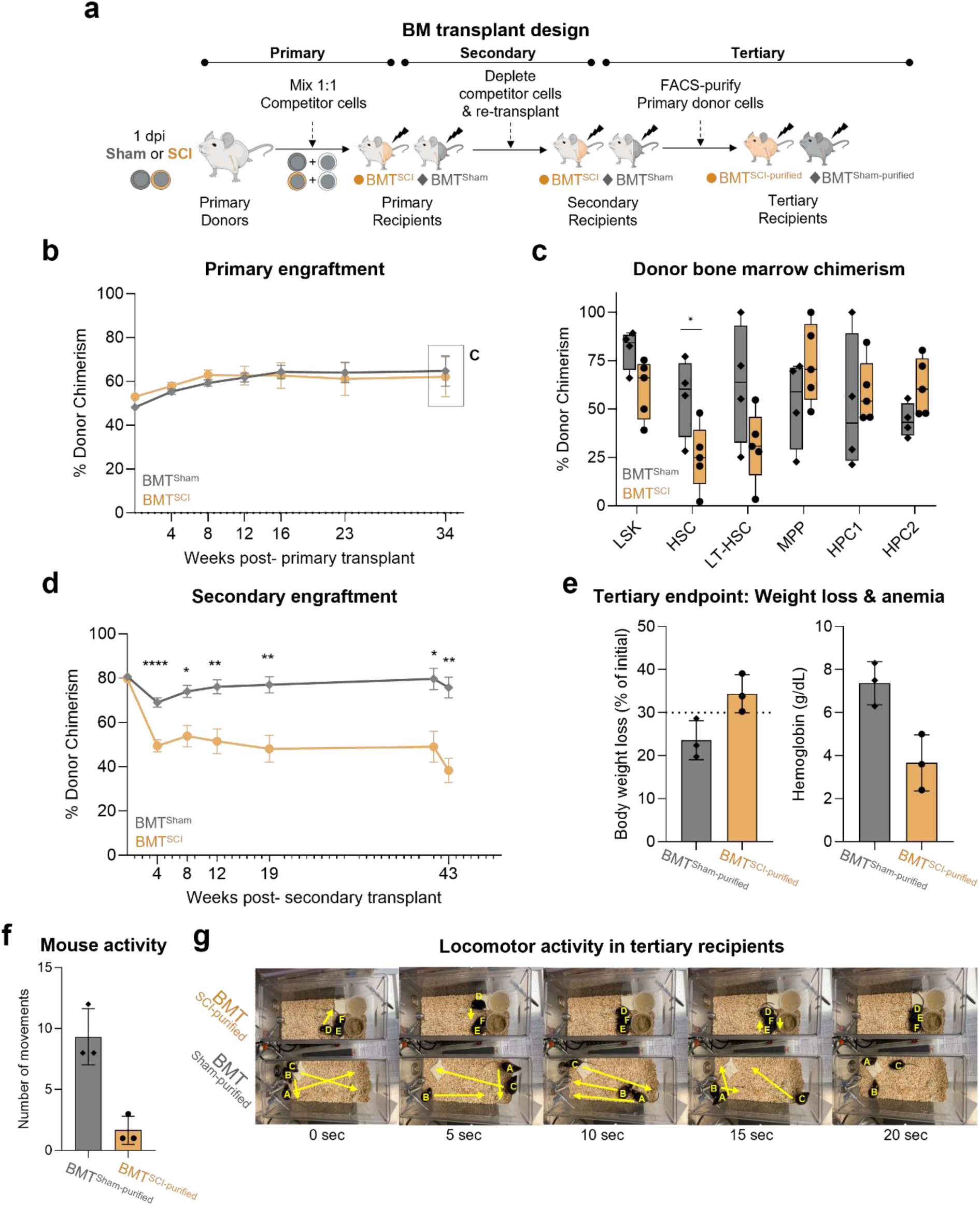
SCI induces long-term HSC defects within 1 dpi. **(a)** Schematic of the primary, secondary, and tertiary bone marrow transplantations (BMT). Donor and competitor cells were distinguished by CD45.2 and CD45.1 antigens respectively. **(b, d)** Donor chimerism in submandibular blood of primary recipients (b) and secondary recipients (d). Values represent the % of donor-derived cells over time. The box represents the timepoint analyzed in (c). **(c)** Donor chimerism in bone marrow LSK cells and subtypes at the primary endpoint. **(e)** Tertiary transplant design and outcomes. Bone marrow from a subset of secondary recipients was FACS-purified for donor-derived CD45.2⁺ cells and transplanted into lethally irradiated tertiary recipients. Data shown at day 16 post-transplant, the latest timepoint at which all SCI-derived transplants remained on study due to meeting early removal criteria (body weight loss and severe anemia). The dotted line represents the 30% early removal criteria threshold per IACUC protocol. These data suggest failure of SCI HSCs to sustain hematopoiesis. **(f)** Number of movements initiated per mouse throughout the video recording (g; screenshots), as a readout of mouse activity. **(g)** Still frames from video recordings immediately prior to euthanasia. Mice receiving SCI-derived bone marrow exhibited pronounced lethargy and reduced locomotor activity. Yellow tracings indicate movement over ∼5-second intervals. See Supplementary Video 1 for full recordings. Overall, these data demonstrate that SCI induces durable, cell-intrinsic defects in HSC function that impair long-term hematopoiesis. **Experimental repetitions**: Primary and secondary transplants (a-d) are from one experiment that was independently repeated at least twice with similar results. Tertiary transplants in (e-g) were done once due to experimental complexity. **Sample sizes:** Mice per group were as follows: n= 4-5 primary donors, n= 4-5 primary recipients (b-c), n= 11-14 secondary recipients (d), n= 7 (d at 43 week engraftment check timepoint), n= 3 tertiary recipients (e-g). **Statistical analysis**: Line plots (b,d) are from the same mice sampled over time. Data are presented as mean ± SEM. Box-and-whisker plots (c) show the interquartile range (25-75%), with the median indicated by the horizontal line. Whiskers span the full data range. Bar plots (e-f) show the mean ± SEM. Individual data points by mouse are overlaid. Statistical comparisons were performed using mixed effects ANOVA with repeated measures (b,d) and Mann-Whitney U tests (e-f). *p<0.05, **p<0.01, ****p<0.0001. **Exclusion:** No mice were excluded from analyses.

SCI-induced HSC defects were sustained in secondary bone marrow transplant recipients (Fig. 3d), a test of long-term HSC function ^40^. Recipients of SCI donor cells displayed significantly reduced peripheral blood chimerism over a 10-month period (Fig. 3d, S3b-g). These data confirm that SCI compromises long-term HSC self-renewal and maintenance while preserving short-term hematopoiesis through unaltered progenitor activity.

To test whether SCI-imprinted HSC defects are lethal, we performed tertiary transplants with purified CD45.2^+^ bone marrow from SCI or sham donors without co-transplanting a survival dose of naïve CD45.1^+^ cells (Fig. 3a “tertiary”, S3h). This approach determines whether SCI-HSCs ultimately lead to hematopoietic failure, a major determinant of reduced lifespan in both SCI and HSC dysfunction ^41^. Purified cell transplants (i.e., from CD45.2^+^ SCI or sham donors without survival doses of naïve CD45.1^+^ cells) eliminate the compensatory effects of competitor cells, revealing HSC defects. Mice receiving purified bone marrow from SCI donor cell recipients developed acute bone marrow failure within 16 days, with severe anemia, >30% loss of body weight, and clinical distress (ruffled fur, microphthalmia, and lethargy; Fig. 3e-g). Subsequent analysis confirmed panleukopenia in blood, bone marrow, and spleen (Fig. S3i-v, Suppl Vid 1). Collectively, these data establish that without supporting bone marrow cells, SCI HSCs are unable to sustain hematopoiesis after serial transplantation. Thus, we have observed that SCI elicits persistent, cell-intrinsic HSC defects that undermine hematopoiesis across the lifespan.

### SCI-induced epigenetic remodeling restricts chromatin accessibility in HSPCs

scRNA-seq analysis revealed distinct transcriptomic profiles in SCI HSPCs, but the regulatory mechanisms underlying these changes remain unclear. Because defects in proliferation, lineage commitment, and long-term repopulation are often epigenetically encoded, we analyzed the HSPC epigenome ^42^. Chromatin accessibility mapping was used to detect lasting epigenetic changes or abnormal regulatory elements that restrict HSPC development and engraftment in the bone marrow niche^43^. Given the similar transcriptional responses to SCI in HSCs and HPCs, we performed bulk ATAC-seq on HSPCs to generate a comprehensive chromatin accessibility map (Fig. S4). Principal component analysis (PCA) revealed that SCI HSPCs possess distinct chromatin accessibility profiles compared to sham and naïve controls (Fig. S4e). We identified 55,068 accessible regions (ARs), predominantly in non-promoter genomic regions (Fig. 4a, inner circle). Differential accessibility analysis detected 809 differentially accessible regions (DARs), comprising 1.47% of ARs, with the majority (484 DARs, ∼60%) exhibiting reduced accessibility in SCI-HSPCs relative to sham (Fig. 4a). Reduced chromatin accessibility predominated across all genomic annotations, including introns, exons, and promoters (Fig. 4a, outer circle). Sham and naïve HSPCs exhibited similar DAR profiles. A heatmap of z-scores for the top 20 DARs (mapped to the nearest gene) illustrates both increased and decreased accessibility in SCI versus sham and naïve HSPCs (Fig. 4b).

**Fig 4.**
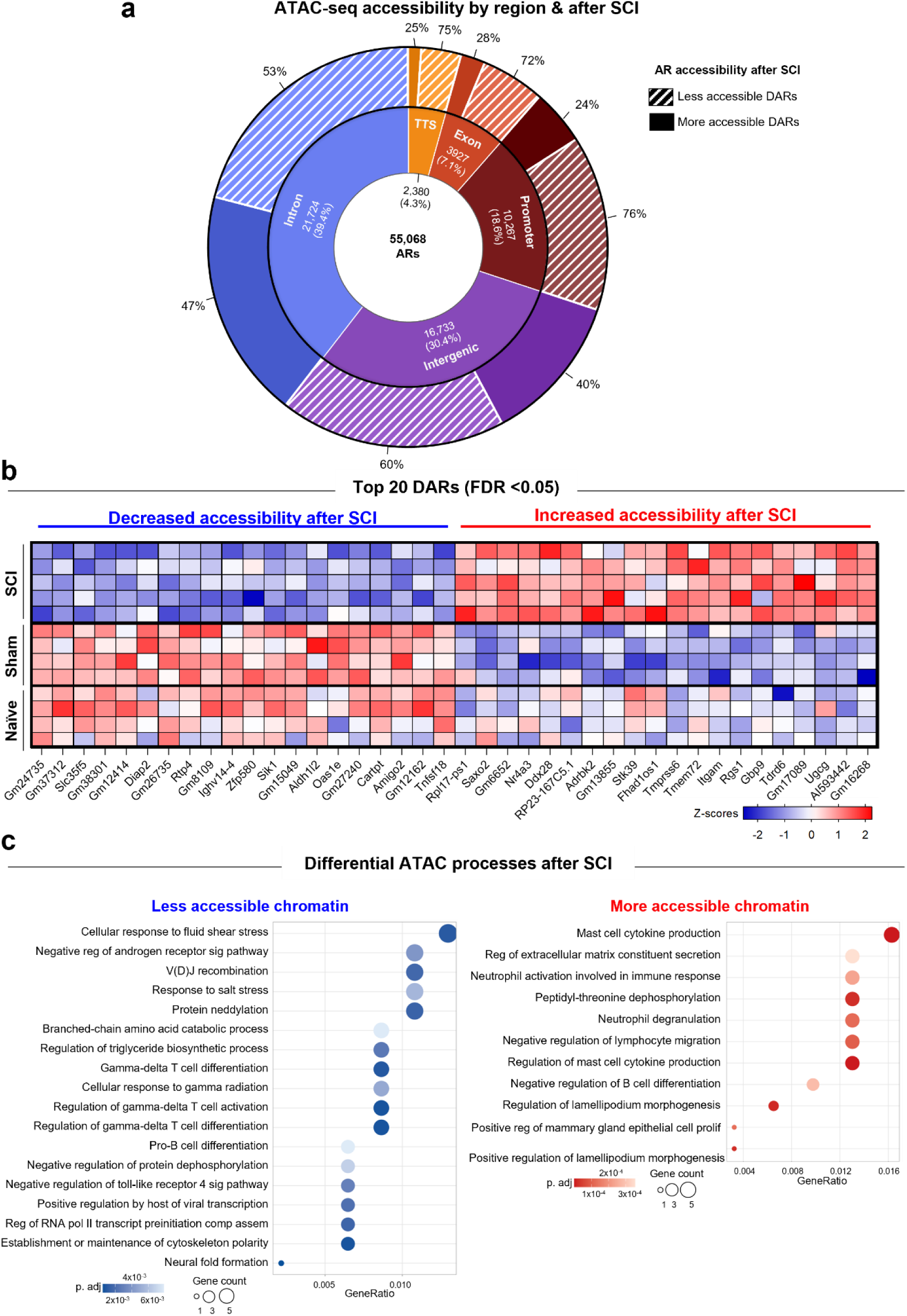
SCI alters the chromatin landscape of bone marrow immune cell precursors within 1 dpi. Bone marrow lineage-depleted, c-Kit enriched mononuclear cells were isolated at 1 dpi from mouse hindlimbs. Cells were obtained from one pooled sample per condition, with each pool comprising bone marrow (n= 5 SCI, n= 4 sham biological replicates). **(a)** Genome annotation of ATAC-seq peaks in the total consensus. Accessible Regions (ARs) were identified (inner ring), prior to assessing SCI-specific effects (i.e., differential accessibility) in outer ring. 809 differentially accessible regions (DARs) identified as FDR <0.05 due to SCI vs. sham (484 less accessible, 325 more accessible). % of all 809 DARs, was split by annotation (i.e., distance from transcription start site; TSS). There were no statistically significant DARs due to sham surgery alone, thus those data are not shown. 37 of 55,068 ARs were unannotated and excluded from the figure. **(b)** Heatmap displaying Z-scores for the top 40 DARs (20 with increased and 20 with decreased accessibility) across individual samples. Each column represents an individual sample (i.e., mouse). Blue indicates decreased accessibility. Red indicates increased accessibility. **(c)** Pathway enrichment visualization for ATAC-seq genes mapped to nearest DARs that are less accessible (blue) and more accessible (red). The GeneRatio reflects the proportion of genes mapping to each pathway. **Exclusion:** One sham mouse was excluded from all ATAC-seq analyses (n= 4 out of 5 included) due to low sample quality upon sequencing.

Pathway analysis of DAR-proximal genes showed that regions with decreased accessibility after SCI were significantly enriched for genes involved in DNA damage response, cytoskeletal organization, transcriptional regulation, and cellular stress responses, including gamma irradiation and heat shock (Fig. 4c, blue). These results indicate that SCI promotes epigenetic silencing of protective and homeostatic programs in HSPCs ^42,44^. In contrast, regions with increased DNA accessibility in SCI HSPCs were enriched for genes associated with immune activation, cytokine signaling, and cell motility (Fig. 4c, red).

### SCI suppresses DNA repair and redox defense programs in HSCs, compromising their resilience to genotoxic stress

To delineate the downstream pathways contributing to HSC dysfunction after SCI, we used our scRNA-seq and ATAC-seq datasets to focus on genes with concordant reductions in expression and promoter accessibility (Fig. S5a). This analysis identified *Fen1* and *Lig1*, which are critical for Okazaki fragment processing and base/nucleotide-excision repair. scRNA-seq further revealed broad SCI-dependent suppression of genes essential for genome integrity and stress response, including *Atm, Brca1, Brca2, Rad51b, Chek1, Fanca, Fanci, Pclaf, Prdx1, Prdx4, Txn1, Txnl1, Park7,* and *Ppia1* (Fig. 5a, S5b) ^13,44,45^.

**Fig 5.**
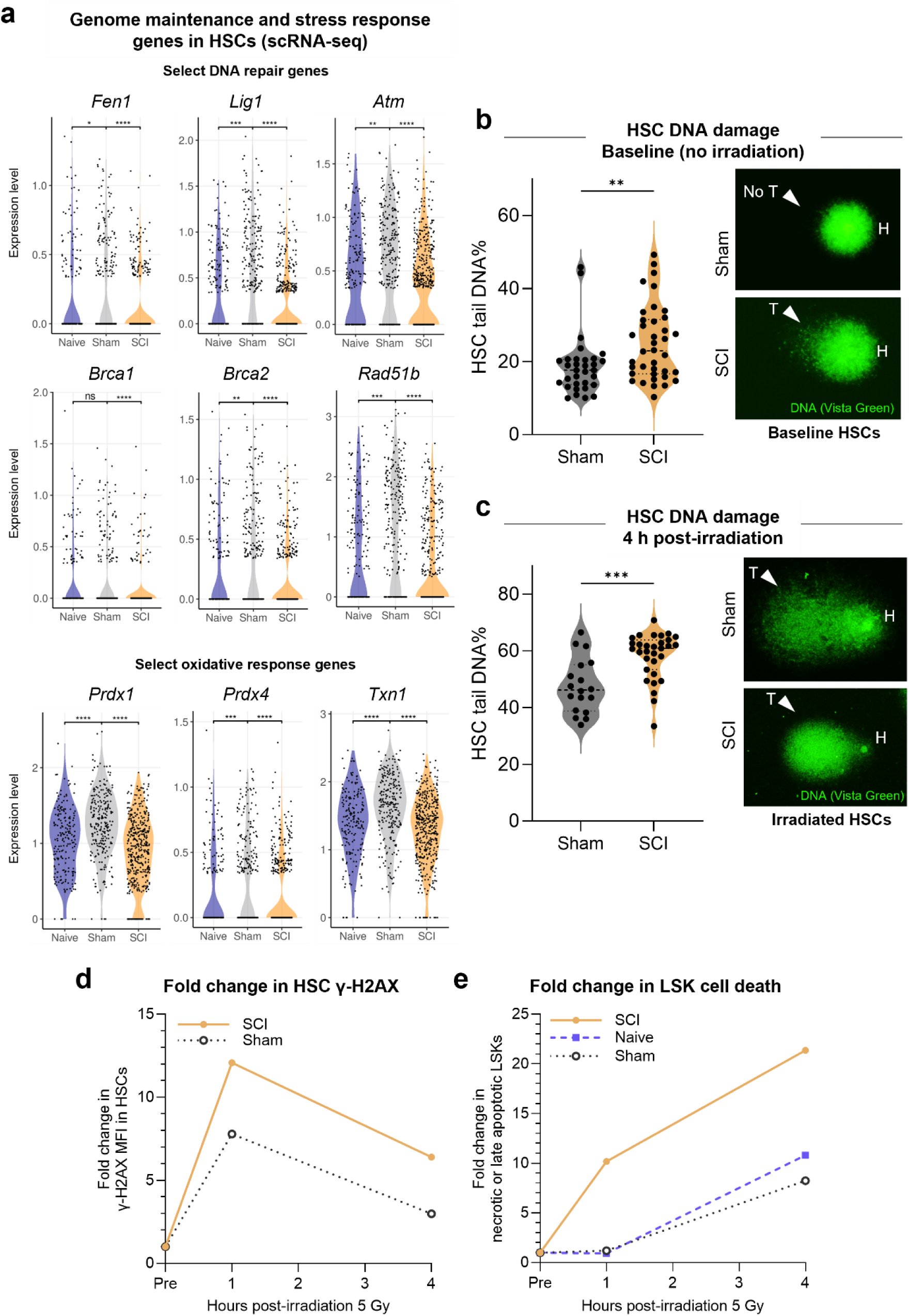
SCI predisposes HSCs to genotoxic damage. Genotoxic damage in SCI HSCs was assessed using transcriptional data and functional HSC assays. **(a)** HSCs were computationally annotated and analyzed in the scRNA-seq dataset (n= 195, 256, and 439 HSCs per group). Violin plots show normalized expression of selected genes involved in genome maintenance and response to cellular stress. Each point represents the expression level of the indicated gene in an individual cell. **(b,c)** Comet assay quantification of DNA damage in FACS-purified HSCs at 1 dpi either at baseline (no irradiation; b) or after irradiation to induce genotoxic stress (10 Gy; c). Data represent the % of DNA found in the tail (i.e., DNA that migrated out of the nucleus), with each point corresponding to a single HSC. Representative comet images. Z-stacks were processed by maximum-intensity projection. DNA was labeled with Vista Green to visualize intact (head; H) and fragmented (tail; T, arrowheads) DNA. Undamaged cells (e.g., sham HSCs in b) show minimal tailing, whereas irradiated HSCs (c) display increased tail formation. Despite genotoxic stress, sham HSCs retain more DNA within the head, indicating less damage compared with SCI HSCs (c, top vs. bottom). **(d,e)** Temporal flow cytometric analysis of γ-H2AX MFI in HSCs (d) or cell death in LSK cells (e) after *ex vivo* irradiation (5 Gy). Time points represent separate aliquots sampled from the same biological preparation; lines connect group means to illustrate temporal trends. Fold changes were calculated by (value at 1 or 4 h)/(pre-irradiation value) with the pre-irradiation value set at 1 for each group and show increased (vs. sham) DNA damage accumulation and persistence at tested timepoints in SCI HSCs. Overall, these data show increased genotoxic damage in SCI HSCs at baseline and after stress. **Experimental repetitions**: Data are representative of two independent experiments (b,c). Data are from one experiment that was independently repeated at least twice with similar results (d,e). **Sample sizes:** Mice per group were as follows: n= 5 (a), n= 9-10 (b-e). Data in (b,c) correspond to n= 32 or 38 HSCs (b) and n= 18 and 31 HSCs (c) analyzed from one pooled sample from n= 9-10 mice to reach sufficient end-gate cell numbers. **Statistical analysis:** Violin plots (a,b) show the median (dotted line, b). Data in (d,e) represent the mean. Statistical comparisons were performed using unpaired two-tailed Wilcoxon rank-sum tests across samples (a) and Mann-Whitney U tests (b,c). *p<0.05, **p<0.01, ***p<0.001, ****p<0.0001. **Exclusion:** No mice were excluded from analyses.

Together, these transcriptional and promoter-accessibility losses implicate broad suppression of DNA damage response and redox defense programs in SCI HSCs (Fig. 5a, S5b). To determine if DNA damage checkpoint activation drives proliferative arrest, we measured baseline DNA damage in freshly isolated HSCs using comet assays ^46^. We found that SCI HSCs exhibited markedly higher endogenous DNA damage than sham and naïve controls (Fig. 5b, S5c), supporting a model in which early DNA damage burden contributes to SCI-induced cell cycle suppression and potential checkpoint activation.

Elevated baseline damage suggests impaired genome maintenance after SCI, but it does not establish whether SCI reduces HSCs’ ability to resolve DNA damage after stress. To test this, we challenged HSCs with irradiation and measured their capacity to limit damage accumulation and preserve genome integrity. Four hours post-irradiation, SCI HSCs sustained significantly greater DNA damage compared to sham controls (Fig. 5c), indicating that SCI increases both endogenous and acute DNA damage. These results demonstrate compromised genome integrity in SCI HSCs underlies defects in homeostatic and stress-induced hematopoiesis.

We next asked whether SCI alters the kinetics of damage resolution and cell death after irradiation in early hematopoietic populations. γ-H2AX was used as a marker of DNA double-strand breaks that resolves as DNA is repaired, enabling assessment of both damage induction and resolution kinetics in HSCs ^47,48^. Genotoxic tolerance was quantified in pooled LSK cells, representing both HSCs and early HPCs following irradiation. We found that γ-H2AX levels were increased in HSCs from both groups, but SCI HSCs showed earlier and prolonged elevation (1 h and 4 h post-irradiation; Fig. 5d), consistent with impaired DNA damage resolution. Over the same period, SCI LSK cells displayed a ∼20-fold increase in late apoptotic/necrotic cells by 4 h, compared to a ∼7-10-fold increase in sham and naïve LSK cells (Fig. 5e).

These findings were confirmed using a lower irradiation dose, allowing assessment at a 21 h endpoint. SCI mice had significantly fewer intact HSCs (ZA⁻ γ-H2AX⁻) relative to their pre-irradiation baseline compared with sham (Fig. S5d-e). The similarity between SCI and naïve at this late timepoint (Fig. S5e) likely reflects distinct underlying mechanisms, with naïve HSCs lacking normal stress-induced priming yet retaining intact repair capacity, whereas SCI HSCs enter irradiation with pre-existing damage and impaired repair, resulting in increased apoptosis compared to naïve controls.

SCI induces inflammatory cytokine production and lipid peroxidation, leading to increased ROS production and DNA damage ^49^. To determine if elevated DNA damage in SCI-HSCs results from impaired ROS attenuation, we assessed their response to hydrogen peroxide (H₂O₂), a potent exogenous oxidant ^50^. We found that sham HSCs exhibited a transient ROS spike followed by a gradual return toward baseline, indicating an intact antioxidant defense (Fig. S6a). In contrast, SCI HSCs failed to resolve ROS, maintaining ∼2-2.5-fold higher levels throughout the evaluation period (Fig. S6b,c).

In summary, SCI induces a transcriptional state characterized by diminished DNA repair and antioxidant capacity, rendering HSCs highly susceptible to genotoxic stress and ROS-mediated damage, which likely contributes to their functional impairment ^48,51^.

### Glucocorticoid receptor blockade reverses SCI-induced HSC dysfunction

SCI imposes a unified repressive state on HSCs, blocking normal stress-induced transcriptional activation, restricting chromatin accessibility at DNA repair and redox defense loci, increasing endogenous DNA damage, reducing tolerance to genotoxic stress, and causing progressive exhaustion of long-term repopulating capacity ^7,52^. These defects indicate that an upstream regulatory signal enforces broad, coordinated suppression of HSC function early after injury. Multiple lines of evidence implicate altered glucocorticoid (GC) signaling as a key upstream mechanism ^53–58^. SCI triggers an immediate surge in circulating GCs via a sympathetic-adrenal neuroendocrine reflex, exposing HSPCs to elevated GC levels during the critical window when transcriptional and epigenetic repression emerges ^59^. GCs, acting through the GC receptor (GR/*NR3C1*), enforce quiescence and restrain G1-to-S phase progression in hematopoietic compartments, mirroring the accumulation of SCI HSCs in G0/G1 and their impaired cycling ^55,60,61^. GR signaling represses transcriptional programs governing cell cycle progression, DNA repair pathway activation, and redox homeostasis, and promotes ROS generation and DNA damage-directly paralleling our observed chromatin closure at repair and antioxidant loci, elevated comet and γ-H2AX signals, and failure of SCI HSCs to resolve oxidative stress ^13,60^. GR also reorganizes promoter and enhancer accessibility in HSPCs, linking the early GC surge to widespread epigenetic compaction at injury-sensitive loci ^62^.

To assess whether GR signaling directly targets regulatory elements with reduced accessibility after SCI, we analyzed promoter-proximal DARs for transcription factor motif scanning (Fig. 6a). Promoter-proximal DARs were enriched among regions with reduced accessibility compared to those with increased accessibility (∼2.1-fold; Fisher’s exact p= 1.13×10⁻³, odds ratio = 2.28). We therefore focused on the 66 promoter-proximal regions with reduced accessibility for motif analysis using JASPAR 2024 (Fig. 6a). Scanning these regions revealed the presence of the *NR3C1* motif (GC receptor; GR, motif ID: MA0113.4; Fig. 6b, top) with eight binding sites, including two high-confidence motif matches within the *Fen1* promoter (Fig. 6b, bottom; p≈ 8.7×10⁻⁵ to 8.4×10⁻⁵). ATAC-seq signal tracks confirmed reduced chromatin accessibility at the *Fen1* promoter in SCI compared with sham animals, overlapping both the annotated DAR and the predicted GR motif (Fig. 6b,c). Consistent with this, accessibility at the *Fen1* promoter was significantly reduced in SCI relative to naïve and sham mice (Fig. 6c).

**Fig 6.**
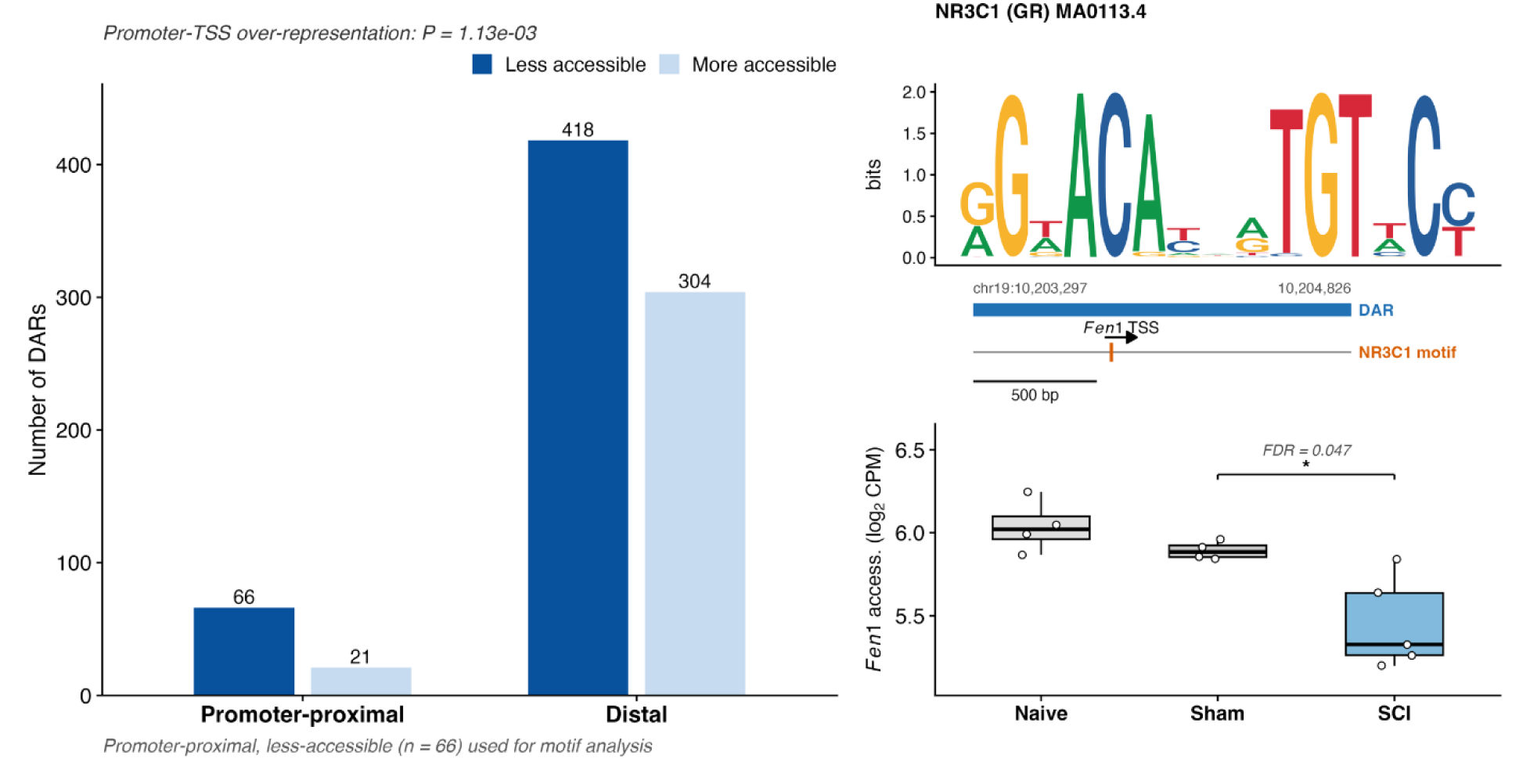
Glucocorticoid receptor motifs are present in promoter-proximal DARs that lose accessibility after SCI. **(a)** Genomic annotation of the 809 differentially accessible regions (DARs; FDR < 0.05) from the spinal cord injury (SCI) versus sham comparison (n= 5 SCI, n= 4 sham biological replicates), stratified by direction of change; less and more accessible denote reduced or increased accessibility in SCI relative to sham. Promoter-TSS regions are 2.1-fold over-represented among less-versus more-accessible DARs (two-sided Fisher’s exact test P= 1.13 × 10^-3^, odds ratio 2.28). The 66 promoter-proximal, less-accessible DARs were retained for transcription factor motif scanning. (**b, top**) *NR3C1* (glucocorticoid receptor; GR) binding motif (JASPAR ID: MA0113.4). (**b, bottom**) schematic of the *Fen1* locus showing the DAR (blue bar; chr19:10,203,297-10,204,826), the transcription start site (arrow) and an *NR3C1* motif match (orange tick; motif scan P= 8.4-8.7 × 10^-5^, both strands). (**c**) *Fen1* DAR accessibility by group (naïve, n= 4; sham, n= 4; SCI, n= 5), shown as log2 counts per million. Boxes show the median and interquartile range, whiskers extend to 1.5 × IQR and points are individual replicates (SCI versus sham FDR= 0.047).

These results demonstrate the presence of GR (*NR3C1*) binding motifs within promoter-proximal regions that lose accessibility after SCI, suggesting that surges in systemic GCs, common in humans and animals after SCI, may drive HSC dysfunction by inducing chromatin remodeling that suppresses DNA repair programs ^59,63,64^. Prior studies show that GR can bind and remodel HSPC promoters (e.g., CXCR4), supporting the plausibility that SCI-elevated GCs target the repair gene promoters identified here ^55,65^. While motif scanning does not establish binding or motif over-representation, the presence of GR motifs at promoters of SCI-repressed repair genes supports their regulatory involvement.

If GR signaling is a primary upstream driver of HSC dysfunction, early GR antagonism should prevent, or reverse functional impairment as measured by serial transplantation. The transplant assay provides a stringent, physiologically integrated evaluation of long-term HSC competence, making it an ideal functional test for this mechanistic hypothesis. To this end, we pharmacologically blocked GR activation using RU486 (mifepristone) administered 6 hours post-injury (Fig. 7a). While RU486 did not alter short-term engraftment kinetics (Fig. S7a,b), consistent with early hematopoiesis being progenitor-driven, it restored long-term chimerism in secondary transplant recipients to sham and naïve levels (Fig. 7b, S7).

**Fig 7.**
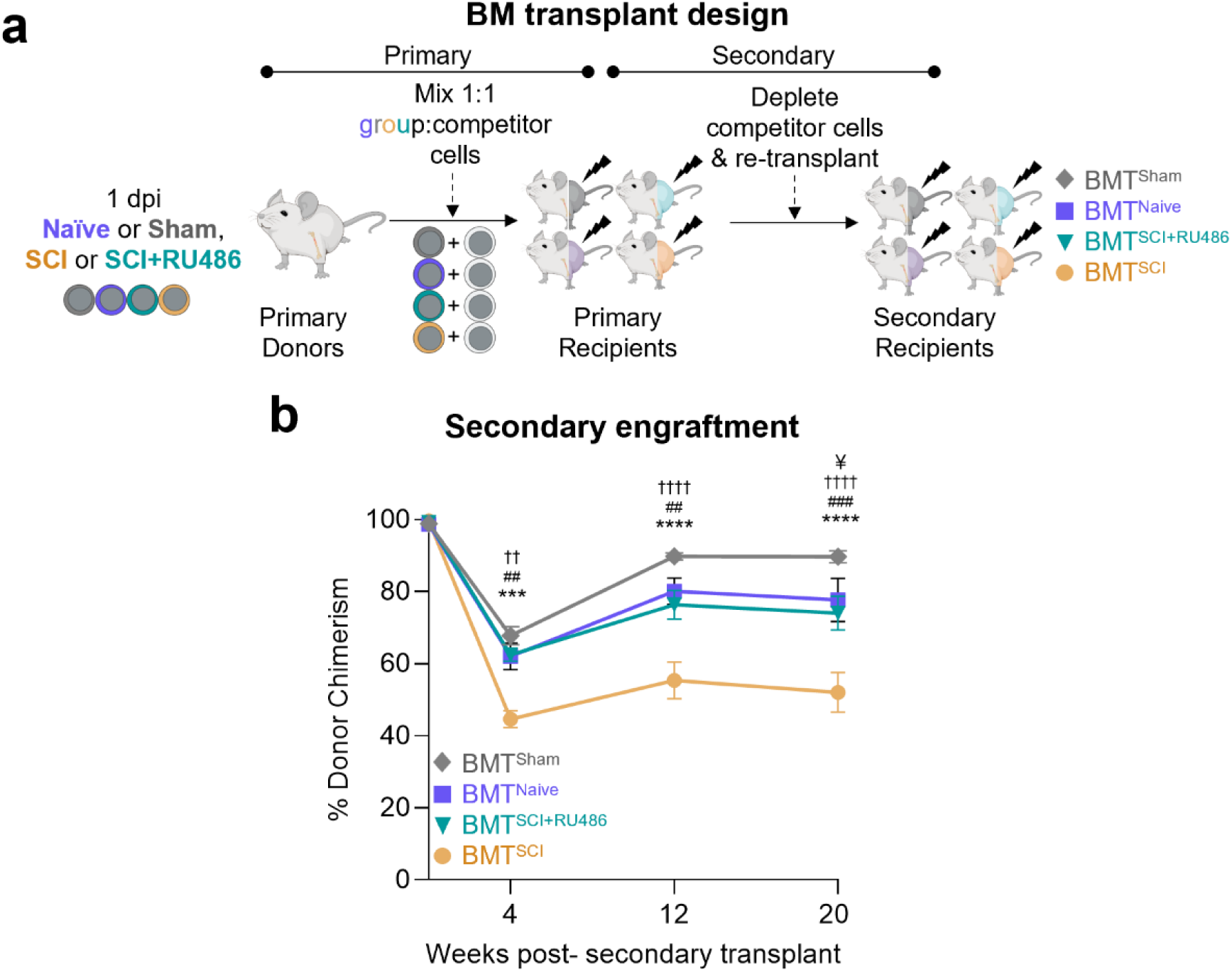
Early stress hormone blockade prevents SCI-induced long-term HSC defects. **(a)** Schematic of the primary and secondary bone marrow transplantation (BMT). Donor and competitor cells were distinguished by CD45.2 and CD45.1 antigens, respectively. SCI mice were randomized such that half received RU486 at 6 hours post-injury (hpi), a clinically relevant intervention timepoint. Naïve mice received anesthesia alone, while sham mice underwent laminectomy without spinal cord damage, modeling surgical stress in the absence of injury. **(b)** Donor chimerism in the submandibular blood of secondary recipients. **Experimental repetitions**: Data are from one experiment that was independently repeated at least twice with similar results, except for the SCI+RU486 group which was included once. **Sample sizes:** Mice per group were as follows: n= 5 primary donors (a), n= 7-8 primary recipients (primary engraftment not shown), and n= 9-10 secondary recipients (b). **Statistical analysis:** Data are represented as mean ± SEM. Statistical comparisons were performed using mixed effects ANOVA with repeated measures (b). †: Naïve v. SCI; #: SCI v. SCI+RU486; *: Sham v. SCI; ¥: Sham v. SCI+RU486. ††p<0.01, ††††p<0.0001. ##p<0.01, ###p<0.001. ¥p<0.05. **Exclusion:** No mice were excluded from analyses.

These findings are important since they demonstrate that SCI-induced HSC dysfunction is not permanent but reflects a GR-dependent repressive state reversible by early GC antagonism. Functional restoration parallels prior evidence that physiological GC signaling primes HSC behavior via *NR3C1* and supports GC signaling as a tractable upstream driver of HSC failure after SCI, thereby revealing a clinically relevant therapeutic window for preserving hematopoietic integrity following neurotrauma ^57^.

## Discussion

Neurotrauma is rarely considered as a modifier of HSC biology; however, our data show that SCI rapidly and durably reprograms HSCs through mechanisms distinct from normal canonical stress hematopoiesis. Instead of activating normal stress responses and proliferative programs to replenish blood cells, SCI enforces a transcriptionally repressed, damage-prone HSC state. This reprogramming emerges within one day of injury and persists through serial transplantation in CNS-intact mice, demonstrating neurogenic imprinting of HSC injury independent of ongoing systemic pathology.

This response is highly directional: sham procedures induce survival, metabolic, and cell-cycle programs, whereas SCI suppresses these adaptive pathways^66^. Early neuroendocrine signaling after SCI fails to activate HSCs and instead imposes a regulatory state that diverges from normal physiological stress responses ^7,9^. Concomitant repression of genome maintenance genes and global chromatin compaction lock HSPCs in a nonproliferative state, consistent with increased DNA damage, unresolved ROS, and loss of HSC function under stress ^13,14^.

Our data illustrate how SCI can remodel HSC identity through epigenetic mechanisms ^42^. SCI consistently closed chromatin at loci associated with DNA repair, cell-cycle regulation, and redox defense (i.e., pathways required for HSC exit from quiescence) ^7,13^. HSPCs, evolutionarily tuned to regulate hematopoiesis under normal stress, are impaired after SCI by a system normally designed to activate them ^7,10,67^. Thus, SCI disrupts normal hematopoietic activation, trapping HSCs in a state primed for inflammatory signaling (evident by upregulation of inflammatory response pathways, response to bacterium, and cytokine production in Fig. 1g), yet unable to meet the demands of producing fully functional mature blood cell types. Decoupling of HSC activation from progenitor cell expansion disturbs normal hematopoietic output driving persistent immune dysfunction known to be present in SCI patients ^21,68^. Notably, the transcriptional suppression in SCI HSCs is accompanied by reduced chromatin accessibility at loci governing DNA repair, cell cycle progression, and oxidative stress responses, hallmarks of a dysfunctional stem cell state ^13,14^. Across transcriptional, epigenetic, metabolic, and functional assays, SCI HSCs exhibit hallmarks of GC overactivation. The ability of early GC receptor (GR) antagonism to restore long-term hematopoiesis strongly suggests that excessive GC signaling is an upstream driver of HSC dysfunction after SCI.

HSCs are essential for sustaining long-term hematopoiesis and immune competence, particularly under physiological stress ^10^. Our data supports that previously unknown HSC defects after SCI likely contributes to reduced lifespan due to impaired hematopoietic output in affected individuals, despite advances in clinical care ^24,69–71^. While our data show that SCI HPCs retain short-term function and can support early hematopoietic recovery, this compensation is not durable. Over time, the inability of SCI HSCs to self-renew and replenish the downstream HPC pool leads to cumulative defects in immune cell production, contributing to chronic immune suppression, anemia, and likely increased infection risk, hallmarks of SCI pathology ^59,72–76^.

Recurrent infections, common after SCI, may also contribute to HSC exhaustion over time ^12,77^. Each infection challenges the system to respond to the hematopoietic demand and then restore homeostasis, stressing the HSC compartment. In healthy individuals, HSCs transiently exit quiescence to replenish HPCs and ultimately produce functionally potent immune cells ^67,78,79^. In contrast, SCI HSCs are transcriptionally and epigenetically suppressed and are unable to effectively mount this response. As HPC reserves deplete (as shown in our transplantation and proliferation data), demand shifts to HSCs, which, due to dysfunction, fail to meet the required hematopoietic needs of the organism. This maladaptive cycle may drive early mortality after SCI through increased susceptibility to sepsis and/or impaired wound healing that occur after SCI ^41,80^. Interventions that preserve HSC function, prevent their dysfunction, or enhance resilience may be critical for improving outcomes after SCI.

Acute SCI triggers a rapid elevation of circulating GCs, a neuroendocrine response that evolved to prioritize survival under extreme stress ^15,59^. While beneficial for short-term adaptation, our data show that, in SCI, excessive GC signaling is maladaptive, silencing hematopoietic programs required for DNA repair and stress resilience. Similar maladaptive effects of the acute post-injury GC surge drive catabolic programs and muscle wasting after SCI ^76^.

In HSPCs, promoters of downregulated DNA repair genes, including *Fen1* and *Lig1*, are enriched for *NR3C1* (GR) motifs, implicating sustained GR activity in establishing the repressed HSC state ^43^. Although motif enrichment does not establish direct GR binding, early GR antagonism (RU486) prevents SCI-induced HSC defects and restores long-term hematopoiesis, identifying GR signaling as a driver of post-neurotrauma hematopoietic dysfunction. Stress hormones also reprioritize metabolism and immunity during acute injury ^81^; therefore, exogenous steroids commonly administered after SCI may further heighten hematopoietic vulnerability by compounding endogenous GC signaling ^82,83^. These findings have immediate translational implications: monitor hematopoiesis early after SCI, when genomic and epigenomic defects are imprinted in HSCs ^19^; target DNA repair and ROS-handling pathways; and preserve bone marrow perfusion, limit oxidative stress, or reverse epigenetic silencing to improve long-term immune competence ^84^. Future studies should define how GCs impair HSC function after SCI (through chromatin remodeling, niche remodeling, selective promoter targeting, or indirect signaling) and test whether selective GR modulation can retain adaptive stress responses while preventing durable hematopoietic failure.

Together, these data identify HSC dysfunction as an early, causal component of SCI-induced bone marrow failure, rather than a downstream consequence of chronic complications. Early GR antagonism preserves long-term HSC function, revealing an early, actionable therapeutic window. This work broadens the framework for understanding SCI pathophysiology and suggests new avenues of therapeutic interventions aimed at protecting hematopoietic integrity in affected individuals.

## Materials & Methods

### Mice & housing

The Institutional Animal Care and Use Committee of the Office of Responsible Research Practices at The Ohio State University approved all animal protocols for this study. All experiments were performed in accordance with the guidelines and regulations of The Ohio State University and outlined in the Guide for the Care and Use of Laboratory Animals from the National Institutes of Health. C57BL/6J (strain #000664; CD45.2) and BoyJ mice (strain #002014; C57BL/6-CD45.1) were purchased from The Jackson Laboratory. Mice were fed commercial food pellets and chlorinated reverse osmosis water ad libitum while housed (≤5/ cage) in ventilated microisolator cages layered with corn cob or alpha-dry soft bedding in a 12 h light-dark cycle at a constant temperature (20 ± 2 °C). Adult male mice (8 weeks or older) were used for all experiments because 78% of SCI occur in males and it removes estrus cycle related confounds ^85^. All mice were euthanized and/or blood collected between Zeitgeber Time 1-5, unless otherwise indicated, to control for circadian changes ^86,87^.

### Surgery & post-surgical care

Mice were assigned to groups using a random number generator. The SCI group received a complete spinal cord transection injury at the third thoracic spinal level after laminectomy of the T3 vertebrae. The surgical sham group (sham) received a T3 laminectomy like the SCI group, but without a SCI. Naïve mice were prepped as done for surgical mice (anesthetized, shaved, and cleaned), but immediately placed back into their cages. Mice were not given dietary supplementation (e.g., high calorie pellets) after surgery to control changes in gut microbiota and/or body fat as a confounding variable. For all surgical procedures, mice were anesthetized with ketamine (120 mg/kg, i.p.) and xylazine (10 mg/kg, i.p.). Mice did not receive prophylactic antibiotics, as these have been shown to alter hematopoiesis ^88^. Mice that survived longer than 1 dpi received daily saline injections s.c. for the first 5 dpi. On the day of euthanasia, bladder expression was omitted to reduce stress-related changes in leukocyte mobilization.

### Blinding & randomization

Mice were randomized at each appropriate step of experiments. On arrival to the animal housing facility all age-, sex-, and strain-matched mice were combined into one bin from different delivery boxes to minimize shipping stress or cohort confound. Mice were distributed to separate cages in succession (ex., 1st mouse into the 1^st^ cage, the 2^nd^ mouse into the 2^nd^ cage., etc) to ensure random distribution of mice per cage (i.e., the ease with which a mouse was selected was not associated with a given cage). Cages were then randomized to groups using a number generator. During sample processing, mice were assigned a number ID without group identifier to prevent sample handling bias. Samples were unblinded during flow cytometric instrument set-up as baseline gates and voltages were set on control groups and ensure the panel worked properly prior to data acquisition. For cell culture assays including the Methocult assay, the researcher plating and counting was blinded to the experimental group. For quantification of CFU, a second identifier on a removable sticker was placed on top of the original label to prevent bias in colony enumeration as often CFU was counted after experiment completion and unblinding for analyses of other outcomes. For transplant assays, the researcher was blinded to groups to prevent unequal treatment throughout the engraftment periods.

### Tissue collection & processing

Mice were terminally anesthetized with ketamine (0.1mL, 100 mg/mL) and xylazine (0.1 mL, 5 mg/mL) i.p. for euthanasia and lack of pain response confirmed.

#### Blood

Approximately 0.3-0.4 mL of blood was collected via right cardiac puncture through a 26 G needle (BD, cat # 309597) into EDTA tubes (Fisher, cat # 22030402) for analysis on a Veterinary Hematology Analyzer (Element HT5, Heska Corp Inc. now Antech Diagnostics) and/or by flow cytometry. Care was taken to obtain the same amount of blood across all mice in each experiment to prevent differences in the level of exsanguination as a hematopoietic stressor. Mice were euthanized around approximately the same time of day between experimental replicates to account for circadian rhythm changes in cell mobilization and function.

#### Bones

Bones (femurs and tibiae) were removed using ethanol-sterilized gloves and scissors/forceps and cleaned of muscle using tissues and placed into conical tubes containing 5 mL of PBS. Under aseptic conditions in a cell culture hood, bones were transferred to a mortar and excess solution vacuumed and immediately replaced with 2-5 mL of PBS+ 2-10% FBS (consistent volume between mice) before being crushed with a pestle. Suspensions were filtered through 40-70 µm cell strainers using a serological pipette and placed on ice for downstream assays.

#### Spleens

Spleens were grossly dissected and adherent pancreatic tissue removed. Spleens were expressed through 40-70 µm cell strainers using the rubber end of 3 or 5 ml syringes and rinsed with 5 mL of PBS+ 2-10% FBS before being placed on ice for downstream assays

### Bone marrow mononuclear cell isolation and depletion of lineage cells

Whole bone marrow was processed into single cell suspensions as above, brought to room temp, and slowly layered over room temp Histopaque 1083 (Sigma, cat # 10831) before centrifugation (1600 rpm, 20 min, brake turned off). The top 2 mL of media containing platelets was discarded and the middle layer containing mononuclear cells (MNCs) transferred into a new 15 mL polystyrene conical tube and brought to 15 mL with PBS + 2% FBS. This was centrifuged (350-400 × g, room temp, 8 min). The pellet was resuspended in PBS + 2% FBS and placed on ice. This was repeated on the pelleted fraction in experiments that required increased MNC yield. 125 µL of a biotinylated lineage cocktail containing the following were added per 100 × 10^6^ cells: anti-TER119 (Fisher, cat # BDB553672), anti-B220 (Fisher, cat # BDB553086), anti-CD11b (Fisher, cat # BDB553309), anti-CD8 (Fisher, cat # BDB553029), anti-Gr1 (Fisher, cat # BDB553125), and anti-CD5 (Fisher, cat #BDB553019). This was incubated for 45 min at 4°C shaking at 40 rpm horizontally and brought to 15 mL with PBS + 2% FBS before centrifugation at 4°C (350 × g, 5 min). Dynabeads Biotin Binder beads (Fisher, cat # 11047) were washed per manufacturer’s instructions and added to the cells at 10 µL of beads per 1×10^6^ cells. Beads and cells were incubated for 30 min at 4°C shaking at 40 rpm horizontally and the volume increased to 7 mL with PBS + 2% FBS. Samples were placed in Dynamag-15 (Fisher, cat # 12301D) for 2 min. The beads were tightened by twisting and the supernatant was removed carefully in aliquots into a new tube. This lineage-depleted MNC sample was then centrifuged at 4°C (350 × g, 8 min) and resuspended in PBS + 2% FBS for flow staining (see appropriate methods section).

### HSPC staining of whole bone marrow for flow cytometry

2-10 × 10^6^ cells were aliquoted per sample or pooled together and distributed across control tubes. A biotinylated lineage antibody cocktail was made combining anti-TER119 (Fisher, cat # BDB553672), anti-B220 (Fisher, cat # BDB553086), anti-CD11b (Fisher, cat # BDB553309), anti-CD8 (Fisher, cat # BDB553029), anti-Gr1 (Fisher, cat # BDB553125), and anti-CD5 (Fisher, cat #BDB553019) and incubating it with cells for 15 min at room temp without shaking in the dark. Tubes were washed with 1 mL of PBS + 2% FBS, centrifuged (300-400 × g, 5 min), and resuspended in 100-200 µL of PBS + 2% FBS. Some cells were purposefully killed using heat or ethanol to create a strong positive control sample for compensation and gating. A mixture of fluorophore-labeled antibodies containing the following were added and incubated with samples for 15 min as before to label LSK subtypes: PerCP Cy5.5 Streptavidin (Biolegend, cat # 4052140, PE anti-mouse Sca-1 (Biolegend, cat # 108107), APC-Cy7 anti-mouse c-Kit (Biolegend, cat # 105826), APC anti-mouse CD150 (Biolegend, cat # 115910), PE-Cy7 CD48 (Biolegend, cat # 103424), with or without FITC anti-mouse CD34 (Fisher, cat # BDB553733), with or without Zombie Aqua Fixable Viability Dye (Biolegend, cat # 423102). For chimeric studies, anti-CD45.1 or CD45.2 antibodies were also added. Cells were washed with 1 mL of PBS + 2% FBS and centrifuged, as before, prior to resuspension in either PBS + 2% FBS or with 1X DAPI (an alternative live/dead staining method; Fisher, cat # H21492). Single color controls were stained using 1-2 µL of the appropriate antibodies per tube. Fluorescence-Minus-One (FMO) controls were created for appropriate color controls. Data were acquired on a BD SORP Fortessa (R647800L6068) supported by The Ohio State University Comprehensive Cancer Center Flow Cytometry Shared Resources Center. Total cell counts were calculated using hemocytometry data.

### Flow cytometry staining of blood

Submandibular blood (1-2 drops) or right cardiac blood (0.2-1 mL) was collected into EDTA-coated tubes (Fisher, cat # 22030402) with collection volume and timing standardized across mice to minimize exsanguination- and circadian-related hematopoietic stress. 100 µL of PBS + 2% FBS was added to these tubes and mixed gently before transferring into 5 mL polystyrene flow tubes (Fisher, cat # 149591A). 10 µL of each sample was combined and equally distributed across control tubes. 1 mL of BD Lysing Buffer (Fisher, cat #BDB555899) was added per tube and antibodies immediately added at 1-2 µL of each: FITC anti-mouse CD4 (Biolegend, cat # 100406), PerCP Cy 5.5 anti-mouse CD8a (Biolegend, cat # 100734), PE-Cy7 anti-mouse Ly6G (Biolegend, cat # 127617) or APC-Cy7 anti-mouse Ly6G (Biolegend, cat # 127624), PE-Cy7 anti-mouse B220 (Biolegend, cat # 103222) or APC-Cy7 anti-mouse B220 (Biolegend, cat # 103224) with or without chimeric antibodies: PE anti-mouse CD45.2 (Biolegend, cat # 109808) and APC anti-mouse CD45.1 (Biolegend, cat # 110714).

### BrdU incorporation assay

Mice received intraperitoneal BrdU injections (50 µg/g body weight) immediately after laminectomy or SCI (t=0 which was just prior to suturing the wounds closed) and at 12 h post-surgery. Lineage-depleted bone marrow MNCs were fixed and permeabilized using BD Cytofix/CytoPerm Buffer (BD, cat # 554714) for 15-30 min on ice in the dark (dark after staining). Following centrifugation (300 × g, 5 min), cells were washed with 2 mL 1x BD Perm/Wash Buffer and resuspended in 250 µL BD CytoPerm Permeabilization Buffer Plus (Invitrogen, cat # 561651) at 50 µL per 1 × 10^6^ cells. After a 10 min incubation on ice, cells were washed again and re-fixed/permeabilized in 500 µL Cytofix/CytoPerm Buffer (100 µL per 1 million cells) for 15-30 min to ensure quality nuclear penetration and fixation. To expose incorporated BrdU, cells were resuspended in 500 µL of 300 µg/mL DNase solution (delivering 150 µg per 5 × 10^6^ cells) and incubated for 1 hour at 37 °C. Cells were then washed and stained with 250 µL BrdU stain (FITC anti-mouse BrdU; Fisher, cat # 501030842) solution per 5 × 10^6^ cells for 20 min at room temp in the dark. After washing, cells were stained with 700 µL DAPI stain solution per 5 million cells for 15 min at room temp in the dark. Final washes were performed as described above, and cells were resuspended in 300 µL 1X Perm/Wash Buffer for acquisition. Values were averaged for sham mice, and each experimental sample (sham or SCI) was divided by this value and multiplied by 100 to get a percentage of sham. This was done to normalize batch effects between all 4 independent studies.

### HSC Ki67 and Hoechst 33342 flow cytometry

Lineage-depleted bone marrow MNCs were stained for LSK cells as described above and then for Ki67 and Hoechst 33342. Cells were fixed and permeabilized as follows using the Foxp3 / Transcription Factor Staining Buffer Set (Fisher, cat # 005523). Briefly, these cells were fixed in 1X Fixation Buffer for 30 min on ice, then permeabilized twice using 1X Permeabilization Buffer. Anti-mouse Ki67 antibody (Biolegend, cat # 652410) was added at 2-5 µL per tube and incubated for 30 min on ice. After washing, cells were stained with Hoechst 33342 (2 µg/mL in PBS + 2% FBS) for 30 min on ice, washed again, and resuspended in Permeabilization Buffer. Samples were stored overnight at 4°C protected from light and analyzed by flow cytometry the following day. Cell cycle phase calling was done in a 3-bin scheme as per the field standard ^78,89^: G0 (Ki67^-^DNA^lo^), G1 (Ki67^+^DNA^lo^), S-G2-M (Ki67^+^DNA^mid/hi^) as shown (Fig. S9).

### ROS flow cytometry assay

Lineage-depleted bone marrow MNCs were pooled per group (to obtain sufficient cell numbers) and stained for LSK cells as above and suspended in 1X (5 µM) Probe Solution. 1X Probe Solution was created by adding 100 µL of 100X Probe Stock to 9.9 mL of PBS + 2% FBS. The 100X (500 µM) Probe Stock was created by dissolving 50 µg of H2-DCFDA (Invitrogen, cat # C6827) into 173 µL of DMSO. 100-200 µL of diluted hydrogen peroxide (H_2_O_2_) was added per tube to induce intracellular ROS and tubes were sampled over time via flow cytometry to assess the temporal resolution of ROS content.

### Flow cytometric gating

Gating schemes are shown (Fig. S8-13).

### Single cell RNA-sequencing

Single cell RNA-sequencing (scRNA-seq) was performed to characterize transcriptional changes in HSPCs following SCI. The bone marrow was harvested from the femurs and tibiae of each mouse and combined in equal numbers to create one pooled sample per group, with each pool comprising bone marrow from n= 5 mice. Density gradient centrifugation was used to remove multinucleated leukocytes and erythrocytes. HSPCs were enriched by magnetic selection for c-Kit⁺ surface expression using the MACS system.

Up to 16,000 c-Kit⁺ cells per group were used as input for single-cell library generation using the 10X Genomics Chromium Next GEM Single Cell 3’ Reagent Kit v3.1 (Dual Index) platform, following the manufacturer’s protocol. Libraries were sequenced on an Illumina NovaSeq6000. Raw BCL files were converted to FASTQ format using Cell Ranger v8.0.0 mkfastq, and alignment to the mm10 mouse genome was performed using Cell Ranger count with default settings, including filtering, barcode counting, and UMI quantification.

Downstream analysis was conducted in R v4.3.3 using Seurat v5^90^. Cells were filtered to exclude those with <1,000 or >9,000 detected features, >25,000 total counts, or >5% mitochondrial reads. Data were log-normalized and integrated using anchor-based canonical correlation analysis (CCA). Cell types were annotated based on canonical marker gene expression. Differential expression analysis was performed using the FindMarkers function (min.pct = 0.1, logfc.threshold = 0.25), and genes with adjusted p-values < 0.05 were considered significant.

Gene set enrichment analysis (GSEA) was performed using the fgsea package [https://www.biorxiv.org/content/10.1101/060012v3] and the MSigDB Gene Ontology Mouse Biological Process reference^91^. Cell cycle phase assignment was conducted using the CellCycleScoring function in Seurat. Violin plots were generated using ggplot2 (RRID:SCR_014601), and statistical comparisons were performed using unpaired two-tailed Wilcoxon rank-sum tests.

### scRNA-seq data availability statement

The scRNA-seq gene expression data FASTQ/H5 files in this publication have been deposited in NCBI’s Gene Expression Omnibus (GEO) and are accessible through GEO Series accession number: GSE317444. The scRNA-sequencing code is available upon request.

### ATAC-seq processing, peak calling, and consensus peak set

Bone marrow lineage-depleted, c-Kit enriched mononuclear cells were isolated as described above at 1 dpi and libraries prepped and sequenced (CD Genomics). Raw ATAC-seq reads were processed using the nf-core/atacseq v2.1.2 pipeline executed via Nextflow v23.10.1 ^92,93^. Briefly, raw read quality was assessed with FastQC v0.11.9, adapters were trimmed with Trim Galore! v0.6.7, and reads were aligned to the GRCm38/mm10 reference genome using BWA-MEM v0.7.17 ^94^. Alignments were sorted and indexed with SAMtools v1.17 ^95^, and duplicate reads were removed using Picard v3.0.0. Reads mapping to mitochondrial DNA or ENCODE blacklisted regions ^96^ were subsequently filtered out. Normalized bigWig tracks were generated for visualization using deepTools v3.5.1 ^97^. Narrow peaks were called with MACS2 v2.2.7.1 (--nomodel --shift −100 --extsize 200 -q 0.05) ^98^, and a consensus peak set was generated across all samples with BEDTools v2.30.0 ^99^. Reads within consensus peaks were quantified using featureCounts (subread v2.0.1)^100^, and peaks were annotated to the nearest transcription start site (TSS) with HOMER v4.11 ^101^. Library complexity and fragment size distributions were assessed using ATAqV (v1.3.1) and deepTools (v3.5.1).

### ATAC-seq differential accessibility and functional analysis

Differential chromatin accessibility was determined in R v4.4.1 using the edgeR package ^102^. The count matrix was filtered for low-abundance peaks, normalized, and fitted to a generalized linear model. Contrasts between experimental groups were performed, and differentially accessible regions (DARs) were defined as those with a false discovery rate (FDR) < 0.05. Principal component analysis (PCA) was performed on variance-stabilizing transformed counts using the DESeq2 package v1.28.0 ^103^ to visualize sample relationships. Functional enrichment analysis of DARs was conducted with the rGREAT R package ^104^. Genomic coordinates of DARs (FDR < 0.05), stratified by directionality, were tested for enrichment against Gene Ontology (GO) databases (Biological Process, Cellular Component, and Molecular Function) ^105^. GO enrichment analysis was performed using clusterProfiler (v4.0) with org.Mm.eg.db annotation ^106^. Benjamini-Hochberg correction was applied with adjusted p-value < 0.05 and q-value < 0.2 thresholds.

### ATAC-seq transcription factor motif analysis

Transcription factor (TF) binding motif scanning was performed using the universalmotif R package with position weight matrices from JASPAR 2024^107^. DAR sequences were extracted from the mm10 genome using BSgenome.Mmusculus.UCSC.mm10. Motif scanning was performed with a stringent p-value threshold of 1 × 10⁻⁴, scanning both DNA strands. Promoter-proximal DARs (within ±1 kb of the nearest TSS by HOMER annotation) that exhibited concordant reductions in both chromatin accessibility (ATAC-seq FDR < 0.05) and transcript expression (scRNA-seq adjusted p < 0.05) were selected for focused motif analysis. The GC receptor (*NR3C1*/GR) motif (JASPAR ID: MA0113.4) was specifically queried based on the a priori hypothesis that SCI-induced systemic GC surges drive GR-mediated chromatin remodeling in HSCs.

### Combined ATAC-seq and scRNA-seq data analysis

To identify genes with concordant chromatin and transcriptional changes, DARs were manually integrated with HSPC and HSC-specific DEGs from parallel scRNA-seq analysis (SCI vs sham, 1 dpi). Genes were classified as having concordant decreased accessibility (ATAC-seq FDR < 0.05, logFC < 0) and decreased expression (scRNA-seq adjusted p < 0.05, log2FC < 0) and promoter-associated DARs were defined as peaks annotated within ±1 kb of transcription start sites by HOMER. Two genes (*Lig1* and *Fen1*) passed both filters and were retained as input sequences for promoter-focused motif scanning.

### Colony-forming unit assay

Colony-forming unit (CFU) assays were performed using MethoCult™ GF M3434 methylcellulose medium (STEMCELL Technologies, cat # 3444), a standard medium that contains erythropoietin and various cytokines to support multilineage hematopoietic colony formation from single HSPCs. Bone marrow cells were plated at a density of 20,000 cells per well in 6-well plates, corresponding to 60,000 cells per 3 mL of MethoCult medium. Cultures were prepared and maintained as previously described ^19^. Colonies were enumerated under an inverted microscope between days 9 and 14 after plating. Colony types were identified based on morphology and size, following manufacturer guidelines.

### DNA comet assay

To assess DNA damage in HSCs whole bone marrow was harvested from 9-10 mice per group and pooled into experimental group samples (number of mice per group is indicated in the appropriate figure legend and/or associated text). Lineage-depleted MNCs were isolated and stained for HSCs, defined as Lin⁻ c-Kit⁺ Sca-1⁺ CD150⁺ CD48⁻, and fluorescence-activated cell sorting (FACS) was performed using a BD FACSAria III at the OSU Flow Cytometry Core Facility. Cells were sorted directly into 100% filtered FBS, washed, and resuspended in PBS prior to assay.

For irradiation experiments, purified HSCs were exposed to X-ray irradiation (RS-2000 Small Animal Irradiator; Rad Source Technologies), at the levels indicated in each figure and/or associated text, and incubated for 4 h at 37°C. DNA damage was assessed using the OxiSelect Comet Assay Kit (Fisher Scientific, cat # NC0588768) and 3-Well Comet Assay Slides (Fisher Scientific, cat # NC0569359). Cells were centrifuged and washed twice with ice-cold PBS lacking Mg²⁺ and Ca²⁺, then resuspended at 1 × 10⁵ cells/mL. Comet slides were pre-warmed to 37°C, and cells were mixed 1:10 (v/v) with Comet Agarose. A volume of 75 µL was added per well.

Slides were incubated horizontally at 4°C in the dark for 15-30 min, and then immersed in pre-chilled lysis buffer (∼25 mL/slide) for 30-60 min. Following lysis, slides were transferred to pre-chilled alkaline unwinding solution (∼25 mL/slide) for 1 h. Electrophoresis was performed in alkaline buffer at 1 V/cm for 30 min, maintaining a current of 300 mA. Slides were rinsed twice in pre-chilled distilled water (5 min each), followed by a 5 min wash in cold 70% ethanol. After drying at 37°C for 15 min, DNA was stained with 100 µL/well of diluted Vista Green DNA dye for 15 min at room temperature.

Imaging was performed using a Nikon AXR Scanning Confocal Microscope with a FITC filter. For each condition, 30-50 cells were imaged per well using an air 20× objective (resolution: 2048 × 2048; line averaging: 4×; zoom factor: 2×; z-stack depth: 30 µm; z-step size: 0.127 µm). Comet tail DNA percentage was quantified using ImageJ software, calculated as tail DNA%= (100*Tail DNA Intensity)/ (Head DNA Intensity) where intensity was the pixel intensity over the ROI drawn around the tail vs. head ^108,109^.

### Irradiation-induced LSK cell death and HSC γ-H2AX assay

To evaluate irradiation-induced cell death and DNA damage, lineage-depleted bone marrow MNCs were isolated from pooled samples of 9-10 mice per group (Fig. 5b-e) or left unpooled to allow for individual biological replicates (Fig. S5d-e) and stained for LSK (Lin⁻ Sca-1⁺ c-Kit⁺) populations as described above. This cell suspension served as the common source material for the irradiation time points. Cells were exposed to 5 Gy (Fig. 5d-e) or 1 Gy (Fig. S5d-e) X-ray irradiation using the RS-2000 Small Animal Irradiator (Rad Source Technologies). Following irradiation, samples were incubated at 37°C while shaking (60 rpm, uncapped). Aliquots were removed from samples at baseline (pre-irradiation) and at various timepoints post-irradiation. Because aliquots were taken at different times during incubation, the exact same cells were not measured across all time points, but all time points originated from the same initial pooled preparation. This design ensured internal consistency while minimizing sample loss during irradiation.

Cell death was assessed with DAPI or 1.5 µL of Zombie Aqua Fixable Viability Dye (BioLegend, cat # 423102) at various timepoints after irradiation (the pre-irradiation death rate serving as control) and incubated for 10 min at room temperature. Samples were washed with PBS and centrifuged at 4°C (400-500 × g, 5 min).

Following viability staining, cells were fixed and permeabilized using the Foxp3/Transcription Factor Staining Buffer Set (Fisher Scientific, cat # 005523). Cells were resuspended in 200 µL of 1X Fixation Buffer and incubated for 30-45 min at 4°C, protected from light. Permeabilization was performed by adding 1 mL of 1X Permeabilization Buffer, followed by centrifugation at 4°C (500 × g, 5 min). This step was repeated once to ensure complete permeabilization. Pellets were resuspended in residual volume and adjusted to 200 µL with 1X Permeabilization Buffer. Cells were kept chilled on ice throughout the entire staining process.

For γ-H2AX detection, cells were stained with anti-H2A.X Phospho (Ser139) antibody (BioLegend, cat # 613404) at 1.5 µL per tube for 30 min on ice. After staining, samples were washed with 1 mL of 1X Permeabilization Buffer and centrifuged (500 × g, 5 min). Final resuspension was performed in 300-500 µL of 1X Permeabilization Buffer. Samples were stored at 4°C, protected from light, and analyzed by flow cytometry within 48 h.

To allow comparison across groups, the pre-irradiation value for each group was normalized to 1 or to each mouse pre-irradiation baseline (Fig. S5d-e) and post-irradiation values were expressed relative to that baseline.

### Bone marrow transplantations

Competitive repopulation assays were performed to assess the functional fitness of HSPCs under experimental conditions relative to a control population. The protocol was adapted from previously published work^19^. Briefly, whole bone marrow was isolated and mixed in equal parts from the experimental group (CD45.2^+^) with naïve non-irradiated congenic BoyJ (CD45.1 ^+^) whole bone marrow. This suspension was injected into lethally irradiated (10 Gy split-dose X-ray irradiation approx. 10-24 h apart) CD45.1 recipients via the lateral tail vein, delivering a total of 4 × 10⁶ cells per mouse in sterile saline without supplements. Peripheral blood was sampled via submandibular veins throughout the engraftment window to assess chimerism via flow cytometry. Chimerism was calculated as CD45.2/ (CD45.1+CD45.2) with or without normalization to the input values.

Secondary bone marrow transplantation assays were performed to specifically evaluate long-term hematopoietic stem cell (LT-HSC) function ^40^. These followed the same protocol as primary transplants, except that CD45.1 competitor cells were depleted from the donor suspension using magnetic bead-based negative selection (Dynabeads system), to enrich CD45.2^+^ donor cells in the injectate. A new cohort of lethally irradiated BoyJ recipients was used for secondary transplantation.

Tertiary transplantation assays were conducted to further assess LT-HSC durability and self-renewal. Donor-derived CD45.2⁺ cells were FACS-purified to >90% purity (confirmed post-sort; data not shown) and injected at a dose of 3.5 × 10⁵ total WBCs per lethally irradiated BoyJ recipient.

### *In vivo* pharmacological glucocorticoid receptor blockade

Circulating GCs rise sharply in both humans and rodents within the first 1-3 days post-injury, and we have previously confirmed this surge in our SCI model ^59,63,64,110^. To pharmacologically block GR activation, RU486 (mifepristone; 50 mg/kg, i.p.) was administered at 6 h post-injury in a 0.15 mL injection using a 25-27 G needle and 1 mL syringe. This dose, higher than that used in non-transplant SCI models to mitigate lymphoid deficiency, was selected as a robust approach to ensure near-complete GR antagonism and minimize subclinical signaling ^21,59,63,73^. RU486 was prepared by dissolving a concentrated stock in DMSO and diluting 1:10 in sesame oil to achieve a final concentration of 10% DMSO. The 6 h therapeutic window was chosen for clinical translatability, as it represents a feasible timeframe for intervention following acute SCI.

### Statistics & outlier exclusion

Data were excluded only in pre-defined cases of significant behavioral deviation following SCI or irradiation (e.g., bladder infection, severe lethargy) or technical errors during sample processing (e.g., flow cytometry antibody staining issues). Data exclusions are disclosed in figure legends and were made in conjunction with statistical outlier detection using Grubb’s test and the ROUT method as needed. Randomization and blinding procedures are detailed in the dedicated section. Group sizes were determined a priori when possible, by analyzing preliminary and published data and/or using G*Power with α0.05 and 80% power. All statistical tests were two-tailed. Appropriate statistics and samples sizes as well as experimental replicates are reported in each figure legend. Individual data points in plots are representative of separate mice or cells and indicated in each figure legend. Data were analyzed using GraphPad Prism software v10.2.1 (GraphPad Software Inc., San Diego, CA) unless otherwise specified. Illustrations were created with BioRender using a lab-paid subscription (biorender.com). Figures were generated in Microsoft PowerPoint.

## Acknowledgements

The authors would like to thank Dr. Christina Marion, Rochelle Diebert, and Ajay Medipally for assistance in caring for the mice throughout the post-transplantation period; Eric Naumann for assistance in tissue harvest in the early stages of project development; Dr. Patrick Collins and Mariam Salem for ATAC-seq library prep and submission for sequencing; Weston Misel for his assistance in tissue isolation and dissection.

## Author Contributions

KAR, AMD, and PGP designed experiments; KAR, KAK, EAA, KAM, JH, RK, ACRD, ZG, and MK performed experiments; KAR and KAK performed bone marrow transplantations; EARG, KAR, and KEM performed scRNA-seq analysis. CW and QM performed ATAC-seq analysis. KAR, AMD, and PGP wrote the manuscript; all authors reviewed and edited the manuscript.

## Supplemental Figures

**Fig S1.**
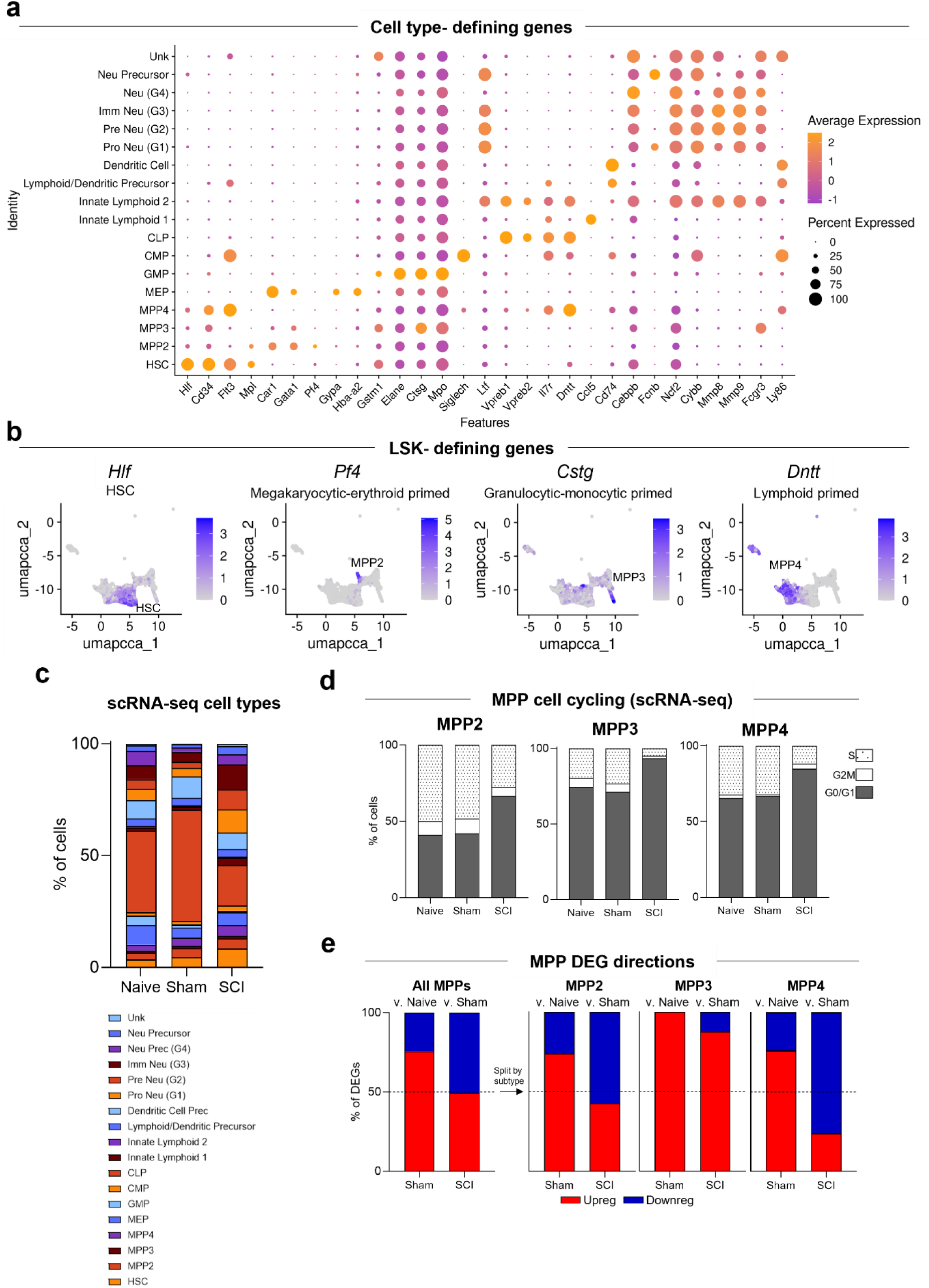
Additional scRNA-seq data in bone marrow immune cell precursors at 1 dpi. **(a)** Dotplot depicting scaled expression of canonical marker genes used for cell type annotation. Data is scaled. **(b)** Feature plots showing expression of canonical LSK marker genes. **(c)** % of each cell type across samples post-QC. **(d)** Cell cycle analysis of the scRNA-seq dataset in non-HSC LSK subtypes at 1 dpi. (dotted: S; white: G2M; grey: G0/G1). Numbers represent the % of cells in each phase. **(e)** % of DEGs in MPP subsets comparing sham vs. naïve (left) and SCI vs. sham (right).

**Fig S2.**
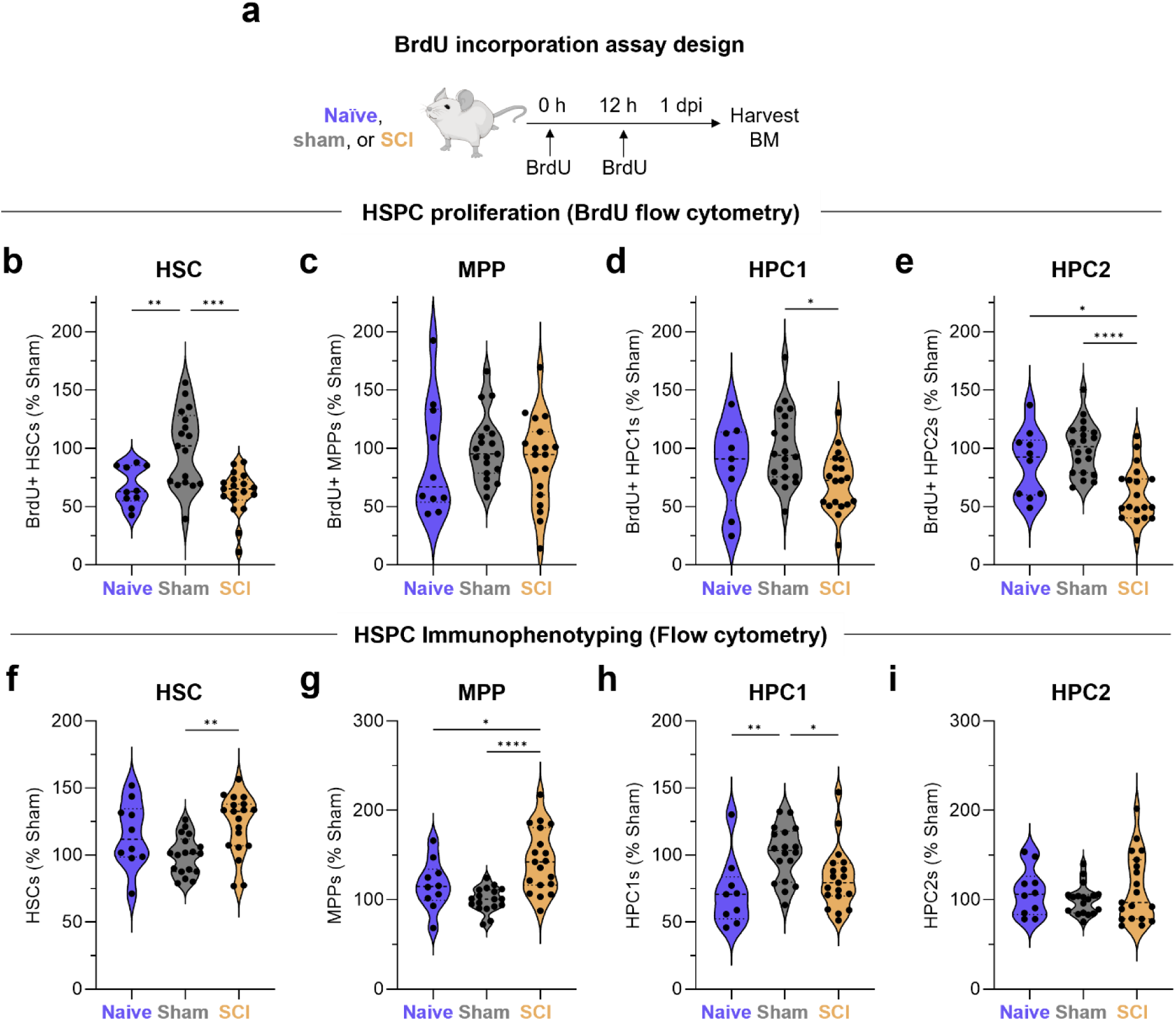
BrdU incorporation assay. **(a)** Schematic of *in vivo* BrdU incorporation to assess HSPC proliferation at 1 dpi by labeling newly synthesized DNA. **(b-e)** Proliferating (BrdU^+^) bone marrow HSCs, MPPs, HPC1s, and HPC2s at 1 dpi, demonstrate reduced proliferation after SCI. **(f-i)** Total numbers of HSCs, MPPs, HPC1s, and HPC2s in bone marrow. **Experimental repetitions**: Data are combined from four independent experiments with their appropriate controls (SCI and sham in two experiments and inclusion of naïve in two additional experiments). **Sample sizes:** n= 10-19 mice per group, with n= 4-5 per group per experiment. **Statistical analysis & normalization:** Values in (b-i) were obtained by back-calculation of flow cytometry percentages to automated hemacytometry counts. These values were averaged for sham mice, and each experimental sample was divided by this value and multiplied by 100 to get a percentage of sham such that values of y= 100 indicate equivalence to sham. This was done to normalize for batch effects between independent studies. Violin plots show the median (dotted line) with individual data points overlaid. Statistical comparisons were performed using ordinary one-way ANOVA (parametric) or Kruskal-Wallis (nonparametric) tests. *p<0.05, **p<0.01, ***p<0.001, ****p<0.0001. **Exclusion:** Two sham mice and one SCI mouse was excluded from analyses due to staining artifacts.

**Fig S3.**
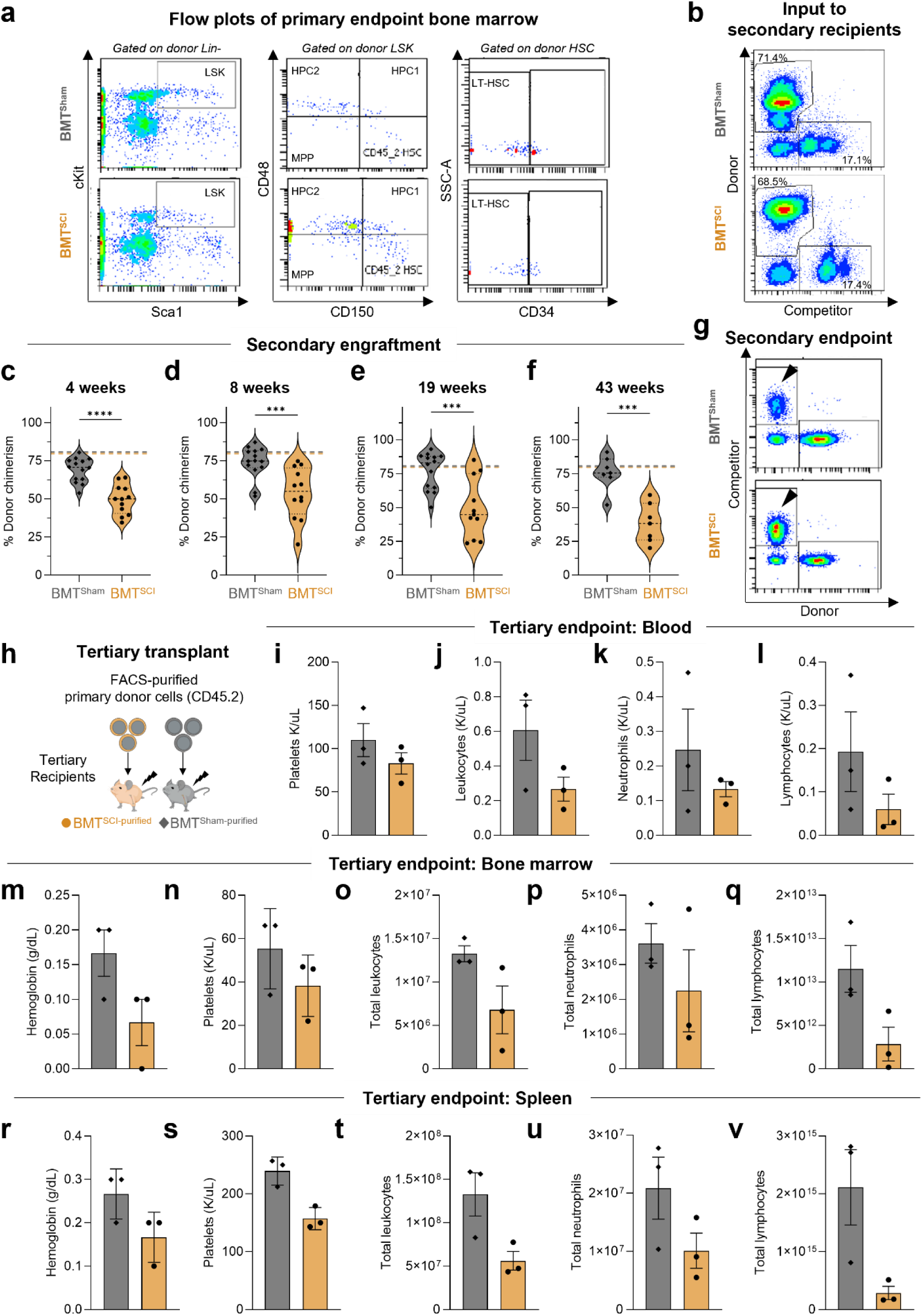
Additional transplant data of sham or SCI bone marrow into naïve mice. **(a)** Representative flow plots corresponding to Fig. 3c. LT-HSCs were gated on CD150^+^CD48^-^ LSK cells and defined by CD34^-^ surface expression. **(b)** Flow plots of bone marrow transplanted into secondary recipients after enrichment for primary donor cells (CD45.2^+^) via magnetic depletion of competitor CD45.1^+^ cells. This enriched bone marrow was transplanted into the secondary recipients shown in Fig. 3a. **(c-f)** Flow cytometry data of peripheral (submandibular) blood from primary recipients showing individual mice from Fig. 3b. Input values plotted as dotted lines (yellow= SCI input, grey= Sham input). **(g)** Representative flow plots corresponding to (f) show increased competitor cells (CD45.1^+^, black arrowheads) in recipients of SCI compared to sham bone marrow, consistent with impaired competitive fitness of SCI-derived hematopoietic cells. **(h)** Schematic of tertiary transplant design. Primary donor CD45.2^+^ bone marrow cells were FACS-purified from secondary recipients. These cells were transplanted into new lethally irradiated BoyJ tertiary recipients. **(i-v)** Automated veterinary hemacytometer data at endpoint in tertiary recipients, collected 16 days after mice met early removal criteria due to >30% weight loss with overt lethargy. **Experimental repetitions**: Primary and secondary transplants (a-g) are from one experiment that was independently repeated at least twice with similar results. Tertiary transplants (h-v) were done once due to experimental complexity. **Sample sizes:** Mice per group were as follows: n= 4-5 primary donors, n= 4-5 primary recipients, n= 11-14 secondary recipients (c-e), n= 7 (f, a randomly selected subset was tested), n= 3 tertiary recipients (i-v). **Statistical analysis**: Violin plots show the median (dotted line) with individual data points overlaid. Statistical comparisons were performed using unpaired t-tests after determining normality and variance (c-f) or Mann-Whitney U tests (i-v). ***p<0.001, ****p<0.0001. **Exclusion:** No mice were excluded from analyses.

**Fig S4.**
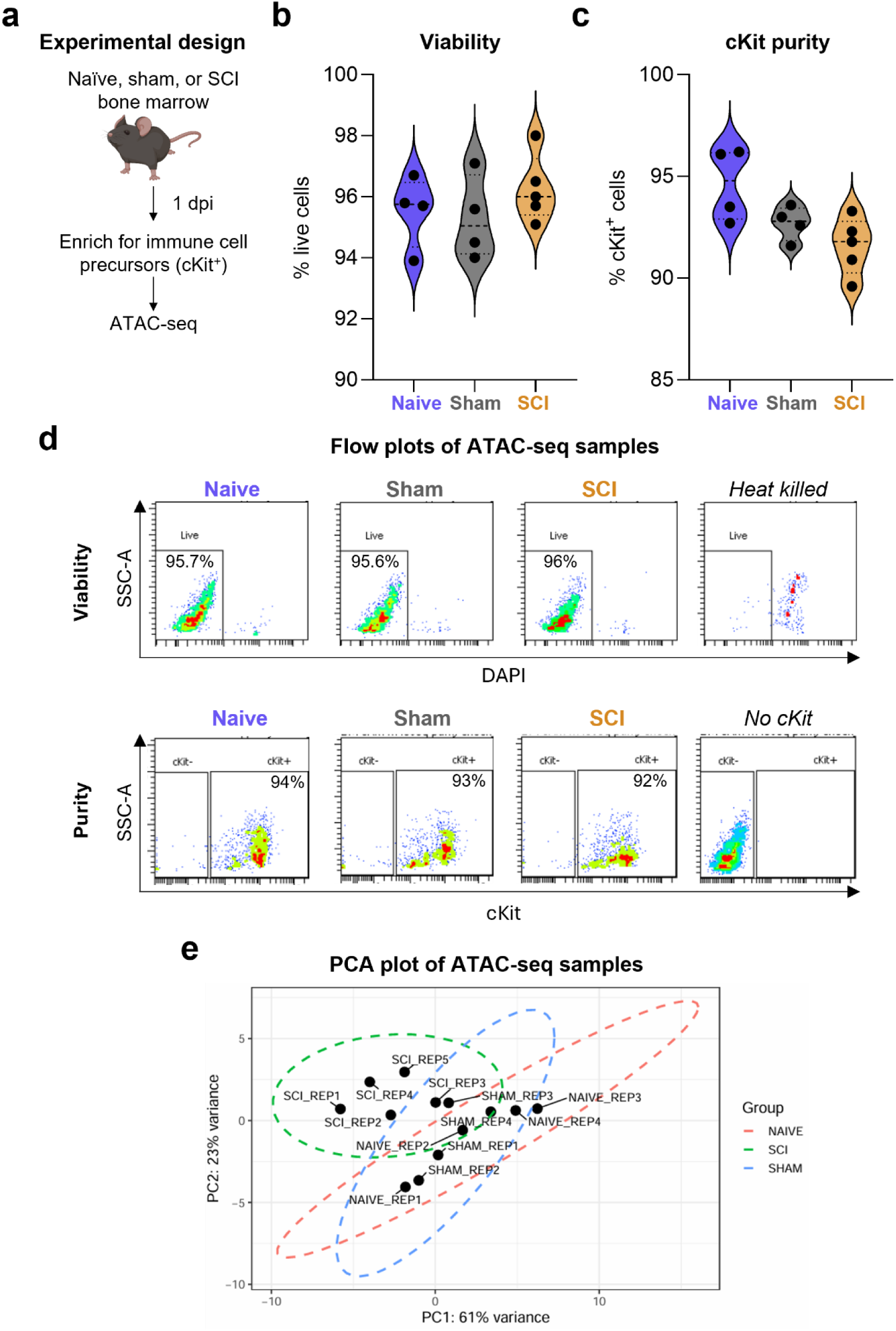
Additional ATAC-seq data in bone marrow immune cell precursors at 1 dpi. **(a)** Schematic of experimental design to assess chromatin accessibility in bone marrow HSPCs at 1 dpi. **(b,c)** Flow cytometry data from an aliquot of samples submitted for ATAC-seq stained for viability using DAPI (b) and purity via c-Kit (c). **(d)** Representative flow plots corresponding to (b,c) gated on singlets (viability plots) or live cells (purity plots). A tube of cells was heated to 55°C to obtain a strong positive control (“heat killed”). A separate tube of cells was not stained for c-Kit to obtain a true negative control (“no c-Kit”). **(e)** PCA of chromatin accessibility profiles in the total consensus peak set. **Statistical analysis**: Each data point represents a biological replicate. Data are presented as mean ± SEM. Statistical comparisons were performed using ordinary one-way ANOVA. **Exclusion:** One sham mouse was excluded from all ATAC-seq analyses (n= 4 out of 5 included) due to low sample quality upon sequencing.

**Fig S5.**
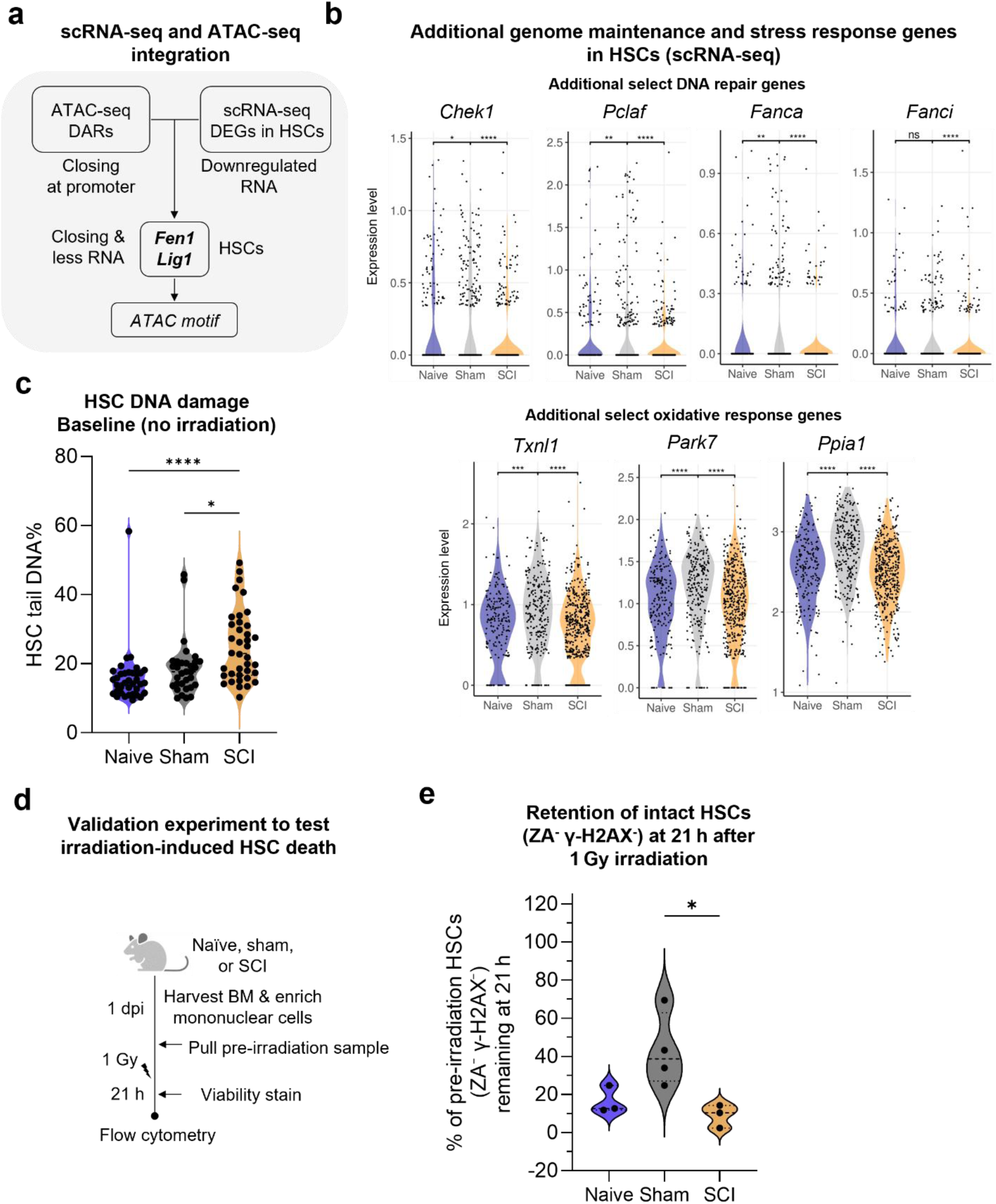
Additional HSC genotoxic stress data. **(a)** Manual integration of downregulated DEGs in HSCs (SCI vs. sham) with DARs from ATAC-seq. Downregulated and less accessible genes (*Fen1* and *Lig1*) were further analyzed using motif analysis to determine transcription factor binding motifs in the promoters. **(b)** HSCs were computationally annotated and analyzed in the scRNA-seq dataset (n= 195, 256, and 439 HSCs per group). Violin plots show normalized expression of selected genes involved in genome maintenance and response to cellular stress. Each point represents gene expression in an individual cell. **(c)** Same dataset as in Fig. 5b, with the addition of cohort-matched naïve mice to demonstrate the absence of significant differences between naïve and sham groups. Individual dots represent HSCs. **(d)** Schematic of the 21 h irradiation validation experiment. Naïve, sham, or SCI bone marrow was harvested at 1 dpi; pre-irradiation aliquots were collected, and mononuclear cells were irradiated (1 Gy, 31 s) and analyzed 21 h later for viability and γ-H2AX. **(e)** Retention of intact HSCs (ZA^-^ γ-H2AX^-^) at 21 h after 1 Gy irradiation, expressed as the % of each mouse’s pre-irradiation HSCs. The y-axis value represents the % of HSCs that survived and resolved genotoxic stress. **Experimental repetitions**: Data are from one experiment that was independently repeated with similar results (c-e). **Sample sizes:** Mice per group were as follows: n= 5 (b), n= 9-10 (c), and n= 3-4 (d-e). Data in (c) correspond to n= 32, 36, or 38 HSCs analyzed from one pooled sample of mice to reach sufficient end-gate cell numbers. **Statistical analysis:** Violin plots (b,c) show the median (dotted line, c). Statistical comparisons were performed using unpaired two-tailed Wilcoxon rank-sum tests (b) and Kruskal-Wallis (nonparametric) tests (c,e). *p<0.05, **p<0.01, ***p<0.001, ****p<0.0001. **Exclusion:** No mice were excluded from analyses.

**Fig S6.**
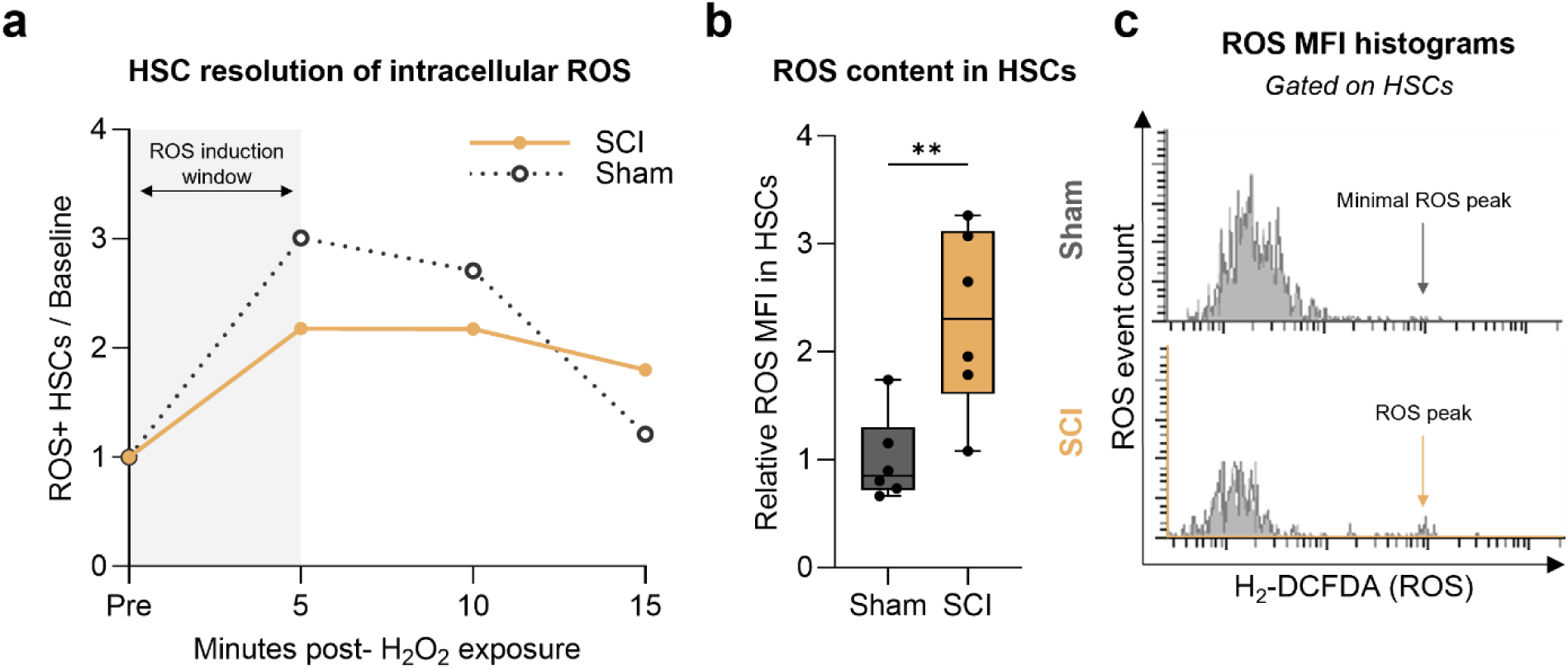
HSC ROS data. **(a)** Resolution of intracellular reactive oxygen species (ROS) in HSCs at 1 dpi following exogenous H₂O₂ exposure, which passively diffuses into cells and increases ROS levels. Changes over time reflect the cell’s ability to reduce ROS through endogenous antioxidant systems. Flow cytometry shows the proportion of ROS^+^ HSCs before and after exposure. Values are normalized to baseline for each group (value of 1= pre-exposure). **(b)** Intracellular ROS content in HSCs (mean fluorescence intensity of H₂-DCFDA) measured at 15 minutes post-H₂O₂ exposure. Values are normalized to the mean sham level (value of 1= equivalent to sham). **(c)** Representative flow histograms. The presence of a peak at the arrow indicates ROS^+^ HSCs. Low counts overall are because HSCs are rare. **Experimental repetitions & sample sizes**: Data are representative of one independent experiment with n= 6 mice per group. Data in (a) are from one pooled sample of mice to reach sufficient end-gate cell numbers. **Statistical analysis:** Box-and-whisker plots (b) show the interquartile range (25-75%), with the median indicated by the horizontal line. Whiskers span the full data range. Statistical comparisons were performed using an unpaired t-test. **p<0.01. **Exclusion:** No mice were excluded from analyses.

**Fig S7.**
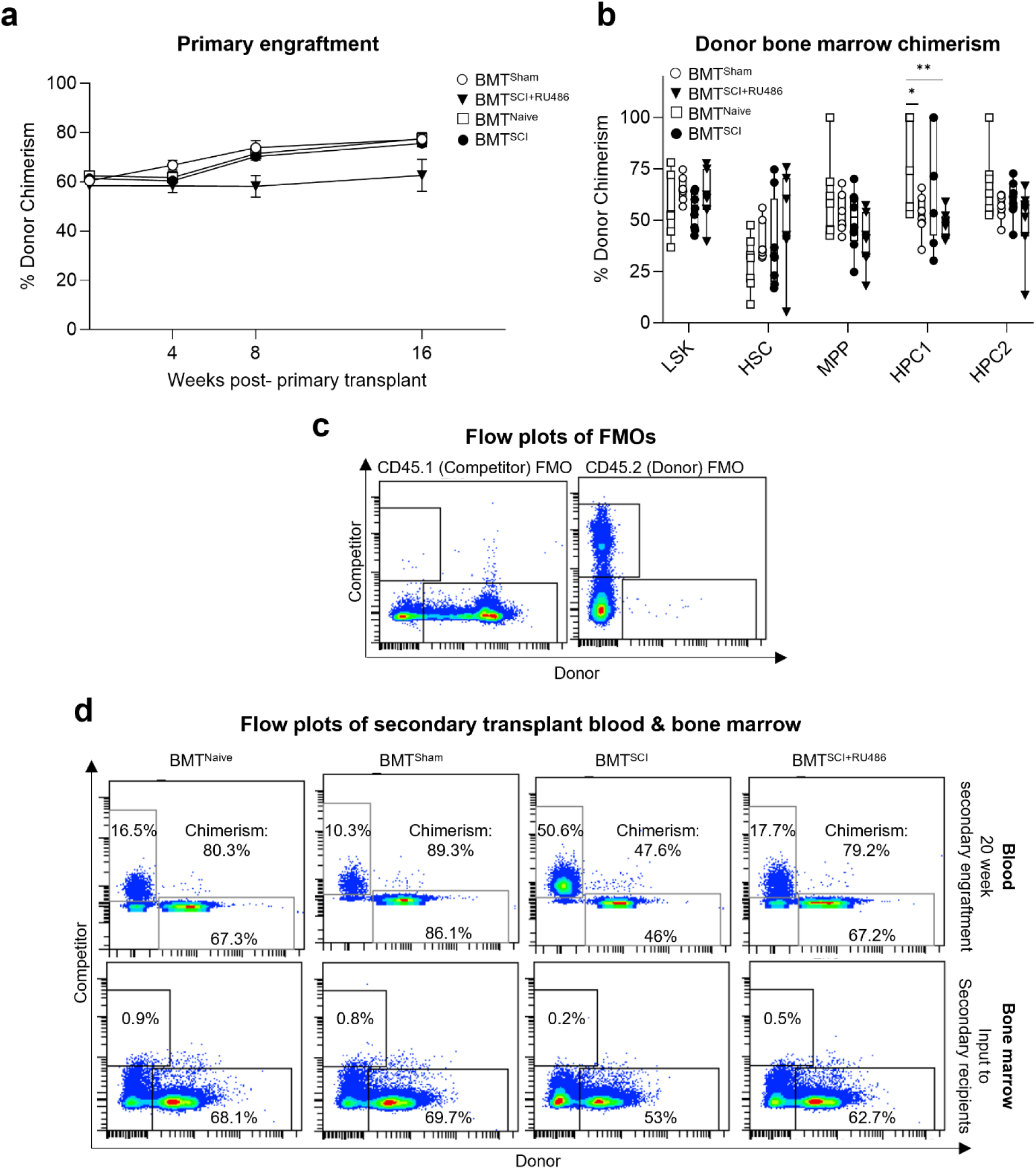
Additional flow plots for data in Fig. 6. **(a)** Flow cytometry data from peripheral blood (submandibular) from primary recipients shown as % donor chimerism. **(b)** Flow cytometry data from bone marrow of primary recipients at endpoint shown as % donor chimerism in LSK cells and subtypes. **(c)** FMO plots validating the gating strategy used to calculate chimerism in transplant experiments. **(d)** Representative flow plots of peripheral blood engraftment at 20 weeks (top) after secondary transplantation or bone marrow (bottom) used for transplantation into these mice 20 weeks earlier to demonstrate the expansion of competitor cells in the SCI group due to intrinsic cell defects. **Experimental repetitions**: Primary and secondary transplants (a-d) are from one experiment that was independently repeated at least twice with similar results. **Sample sizes:** Mice per group were as follows: n= 4-5 primary donors, n= 4-5 primary recipients (b), n= 11-14 secondary recipients (d). **Statistical analysis**: The line plot in (a) contains repeated measurements from the same mice over time; data are presented as mean ± SEM. Box-and-whisker plots (b) show the interquartile range (25-75%), with the median indicated by the horizontal line. Whiskers span the full data range. Individual data points by mouse are overlaid. Statistical comparisons were performed using mixed effects ANOVA with repeated measures (a) and ordinary one-way ANOVA within cell subtypes (b). *p<0.05, **p<0.01. **Exclusion:** No mice were excluded from analyses.

**Fig S8.**
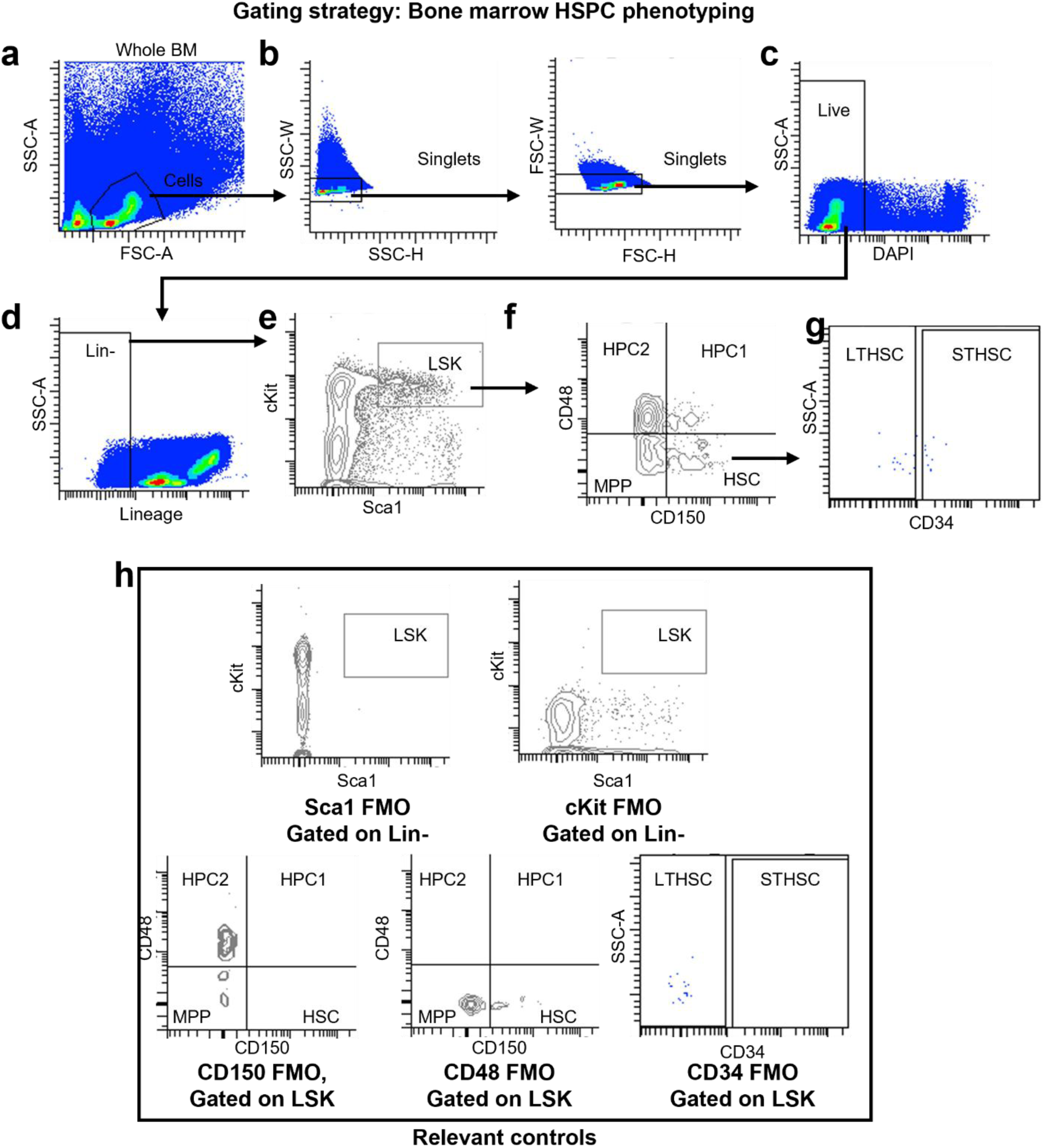
Gating strategy for bone marrow HSPC phenotyping. **(a)** Total bone marrow cells were first identified using FSC-A versus SSC-A to exclude debris. **(b)** Singlet discrimination was performed using SSC-W vs. SSC-H followed by FSC-W vs. FSC-H to remove doublets. **(c)** Live cells were gated by excluding DAPI⁺ events. **(d)** Lineage-negative (Lin⁻) cells were selected using a lineage marker cocktail to remove differentiated hematopoietic cells. **(e)** Lin⁻ cells were then gated on Sca-1 and c-Kit to define the LSK (Lineage⁻ Sca-1⁺ c-Kit⁺) population. **(f)** LSK subsets were resolved using CD48 and CD150 to distinguish HSC, MPP, HPC1, and HPC2 populations. **(g)** Long-term and short-term HSCs were further separated based on CD34 expression. **(h)** Fluorescence-minus-one (FMO) controls for Sca-1, c-Kit, CD150, CD48, and CD34 were applied to the appropriate parent populations (Lin⁻ or LSK) to establish positive gating thresholds and validate marker specificity across all HSPC subsets.

**Fig S9.**
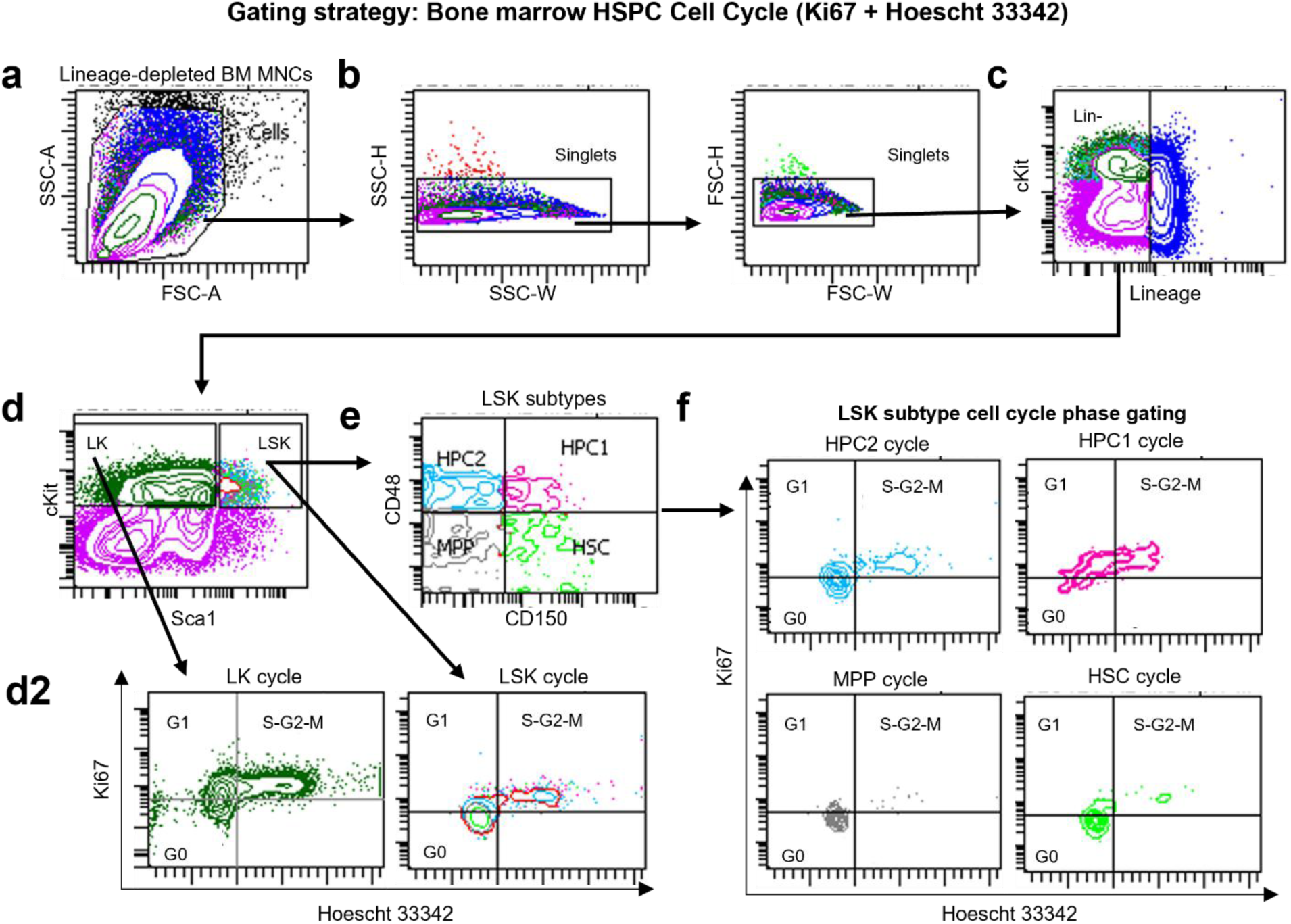
Gating strategy for bone marrow HSPC cell-cycle analysis (Ki67 + Hoechst 33342) **(a)** Lineage-depleted bone marrow mononuclear cells (MNCs) were isolated and identified based on FSC-A and SSC-A to exclude debris. **(b)** Singlet discrimination was performed using SSC-W vs. SSC-H followed by FSC-W vs. FSC-H to eliminate doublets. **(c)** Lin⁻ cells were selected using a lineage marker cocktail. **(d)** Lin⁻ cells were then gated on Sca-1 and c-Kit to identify LSK and LK fractions. **(d2)** Ki67 and Hoechst 33342 intensity were used to define G0 (Ki67^-^DNA^lo^), G1 (Ki67^+^DNA^lo^), and S-G2-M (Ki67^+^DNA^mid/hi^) phases within the LK and LSK gates. **(e)** LSK subsets were resolved using CD48 and CD150 to distinguish HSC, MPP, HPC1, and HPC2 populations. **(f)** Cell-cycle phase distribution (G0, G1, S-G2-M) was quantified within each LSK subtype using Ki67 expression and Hoechst 33342 DNA content. This gating strategy was applied uniformly across samples to ensure consistent identification of cell cycle states within HSPC compartments. **Colors:** HSCs (Lime), grey (MPPs), HPC1s (pink), HPC2s (light blue), forest green (LK cells),

**Fig S10.**
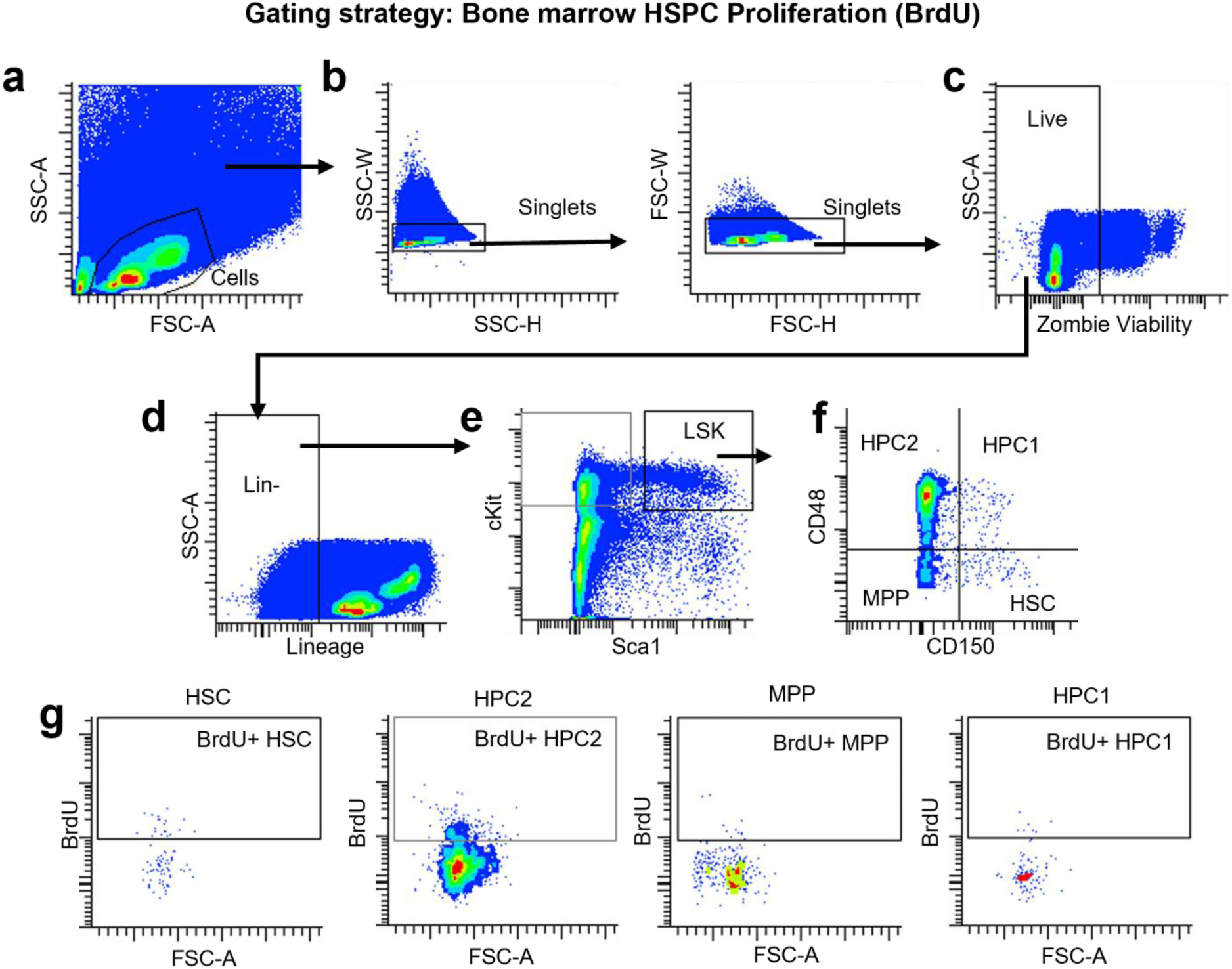
Gating strategy for bone marrow HSPC proliferation (BrdU) **(a)** Total bone marrow cells were identified using FSC-A versus SSC-A to exclude debris. **(b)** Singlets were selected by sequential SSC-W vs. SSC-H and FSC-W vs. FSC-H gating to remove doublets. **(c)** Live cells were gated based on exclusion of Zombie viability dye-positive events. **(d)** Lineage-negative (Lin⁻) cells were selected using a lineage marker cocktail to eliminate differentiated hematopoietic cells, and **(e)** Lin⁻ cells were subsequently gated on Sca-1 and c-Kit to identify LSK cells. **(f)** LSK subsets were resolved using CD48 and CD150 to distinguish HSC, MPP, HPC1, and HPC2 populations. **(g)** BrdU incorporation was assessed within each LSK subtype, with BrdU⁺ fractions identified based on BrdU fluorescence intensity.

**Fig S11.**
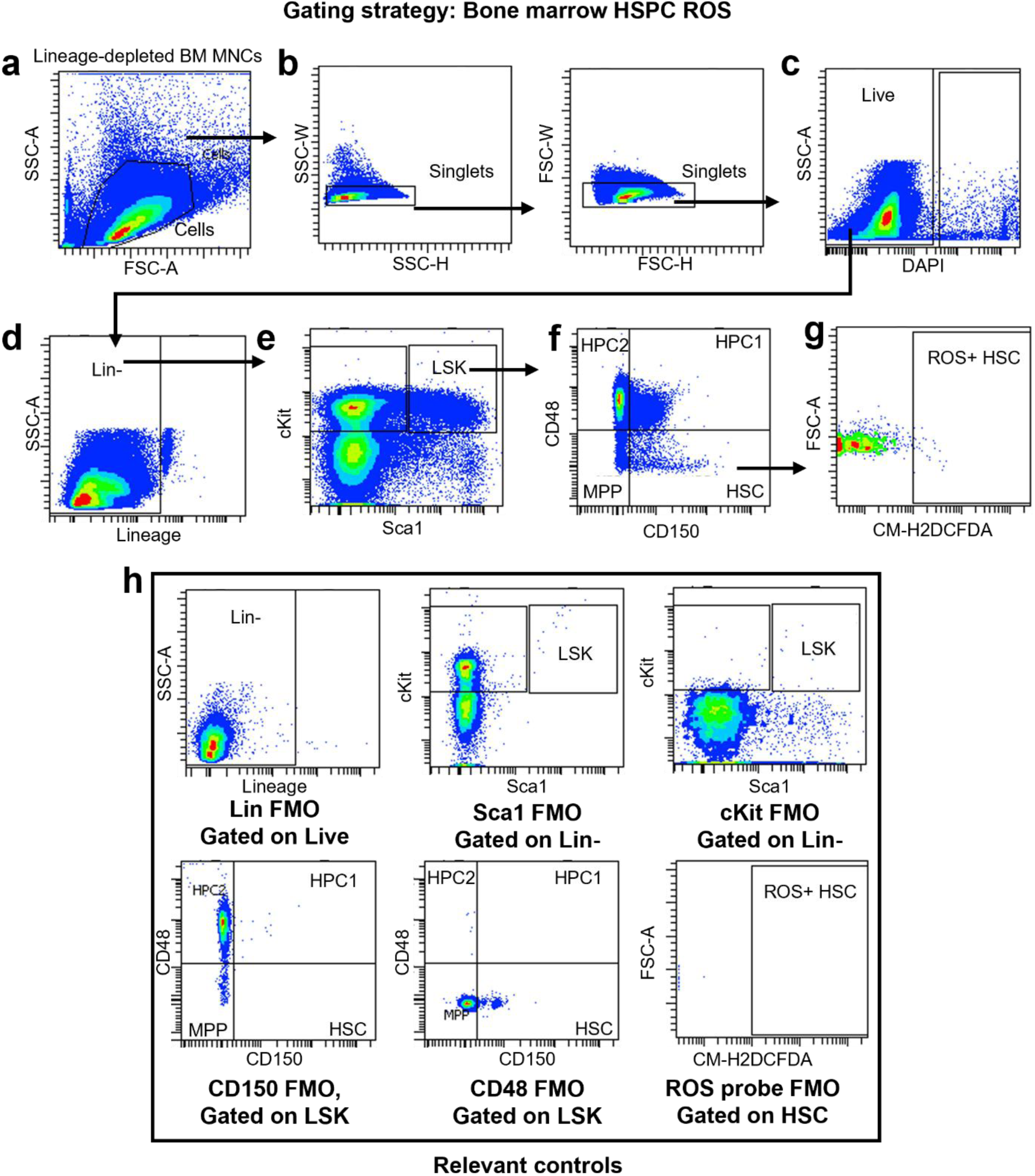
Gating strategy for bone marrow HSPC ROS analysis. **(a)** Lineage-depleted bone marrow mononuclear cells (MNCs) were isolated and identified based on FSC-A and SSC-A to exclude debris. **(b)** Singlet discrimination was performed using SSC-W vs. SSC-H followed by FSC-W vs. FSC-H to remove doublets. **(c)** Live cells were selected by excluding DAPI⁺ events. **(d)** Lineage-negative (Lin⁻) cells were gated using a lineage marker cocktail to remove differentiated hematopoietic cells. **(e)** Lin⁻ cells were subsequently gated on Sca-1 and c-Kit to define the LSK compartment. **(f)** LSK subsets were resolved using CD48 and CD150 to distinguish HSC, MPP, HPC1, and HPC2 populations. **(g)** Intracellular ROS-containing HSCs were identified based on fluorescence intensity of the ROS probe. **(h)** Fluorescence-minus-one (FMO) controls for lineage markers, Sca-1, c-Kit, CD150, CD48, and the CM-H2DCFDA probe were used at the appropriate parent gates (Live, Lin⁻, or LSK) to establish positive gating thresholds and ensure specificity for ROS detection within HSCs and LSK subsets.

**Fig S12.**
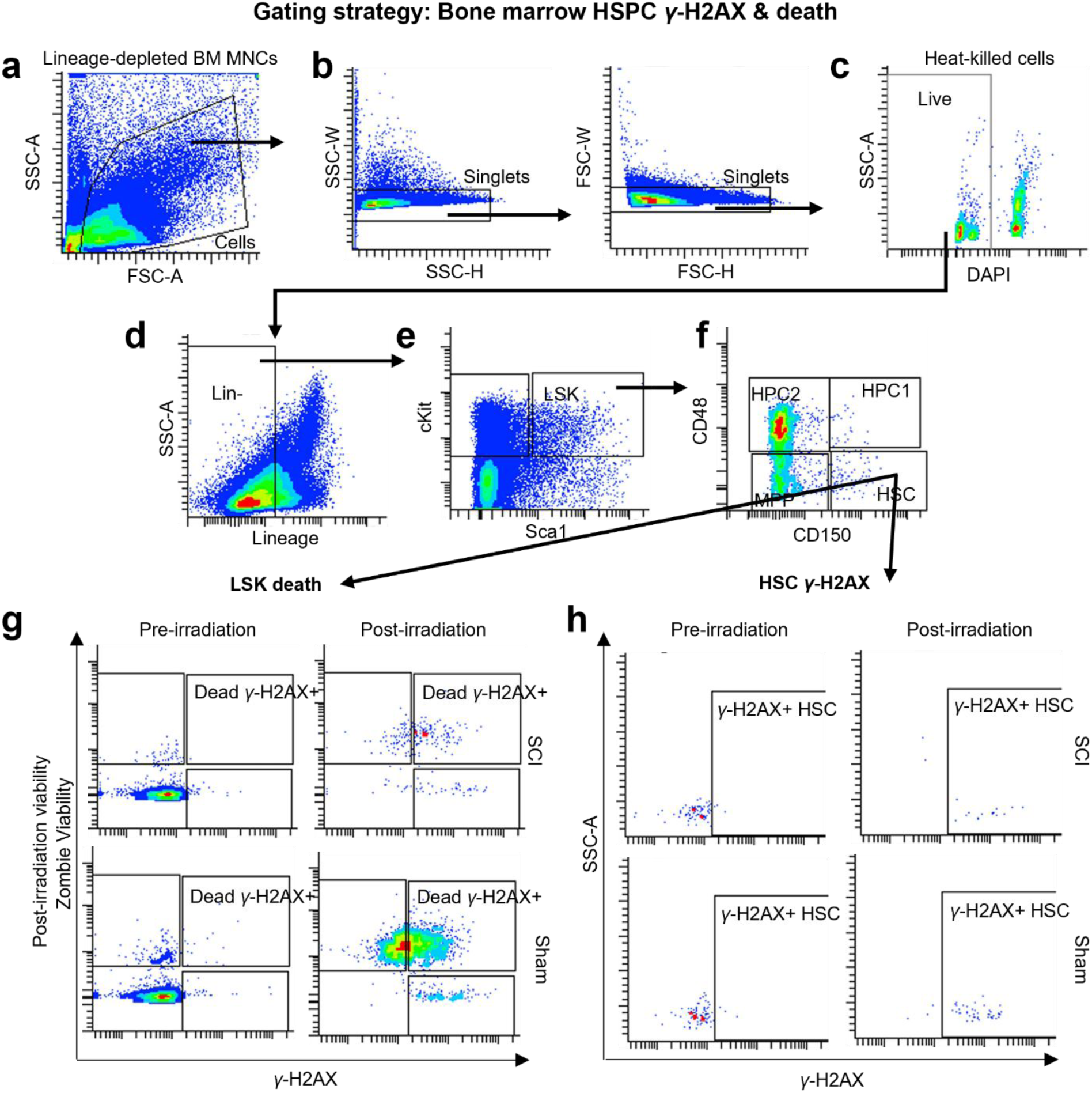
Gating strategy for bone marrow HSPC γ-H2AX and cell death analysis. **(a)** Lineage-depleted bone marrow mononuclear cells (MNCs) were isolated and identified based on FSC-A and SSC-A to exclude debris. **(b)** Singlets were selected by sequential SSC-W vs. SSC-H and FSC-W vs. FSC-H gating to remove doublets. **(c)** Live cells were gated by excluding DAPI⁺ events, with heat-killed cells shown as a positive control for dead-cell staining. **(d)** Lineage-negative (Lin⁻) cells were selected to dump differentiated hematopoietic cells. **(e)** Lin⁻ cells were then gated on Sca-1 and c-Kit to define the LSK compartment. **(f)** LSK subsets were resolved using CD48 and CD150 to distinguish HSC, MPP, HPC1, and HPC2 populations. **(g)** Cell death and DNA damage in LSK cells were assessed by plotting Zombie viability dye versus γ-H2AX before and after irradiation, allowing identification of dead γ-H2AX⁺ fractions. **(h)** γ-H2AX⁺ HSCs were further visualized separately pre- and post-irradiation to illustrate irradiation-induced DNA damage responses. This gating workflow was applied across all samples.

**Fig S13.**
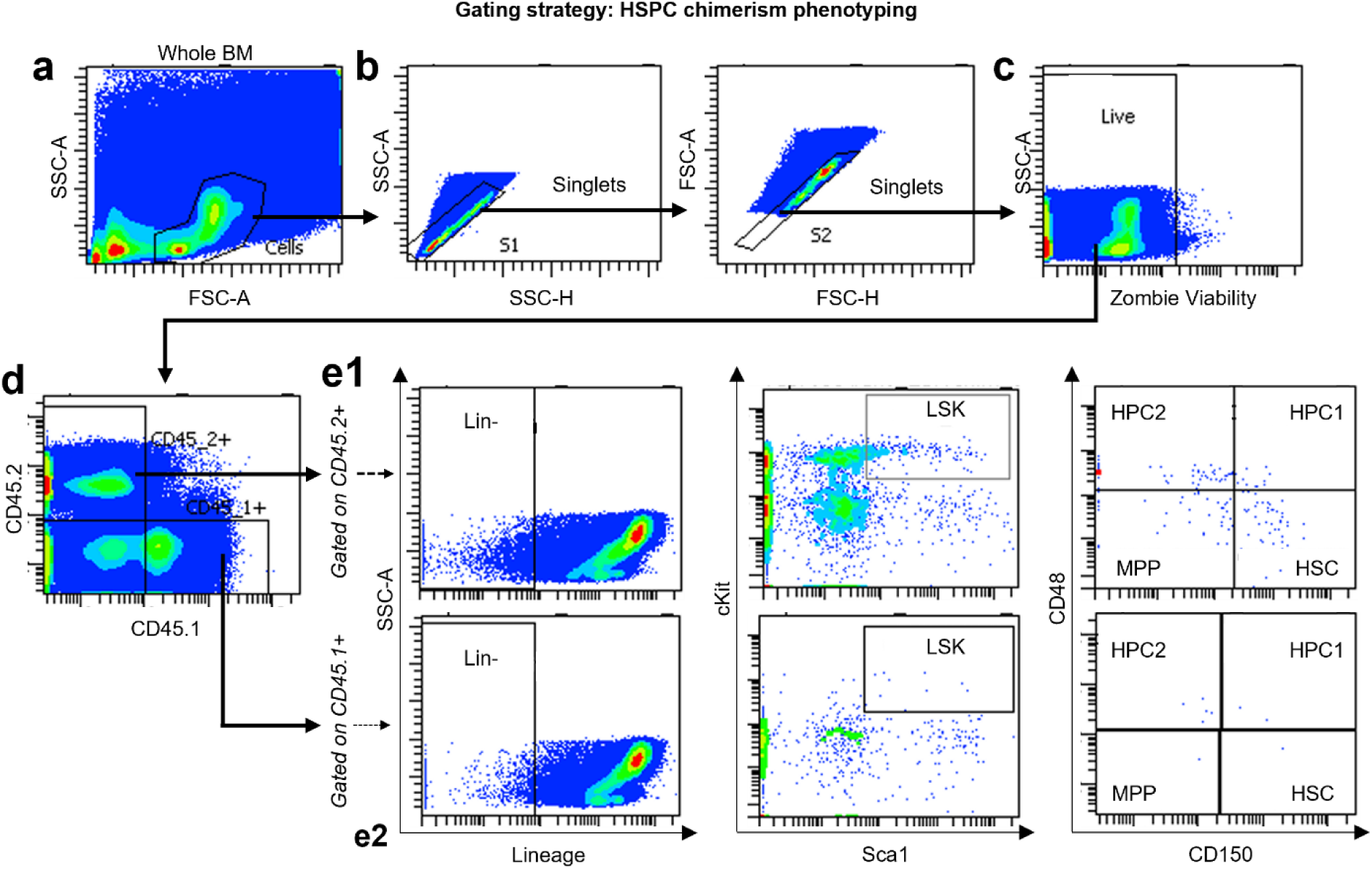
Gating strategy for HSPC chimerism phenotyping. **(a)** Whole bone marrow cells were identified using FSC-A versus SSC-A to exclude debris. **(b)** Singlets were selected by sequential SSC-W vs. SSC-H and FSC-W vs. FSC-H gating to remove doublets. **(c)** Live cells were gated excluding ZA^+^ events. **(d)** Donor-derived (CD45.2⁺) and competitor-derived (CD45.1⁺) cells were distinguished by CD45.1/CD45.2 expression, and downstream gating was performed separately for each fraction. **(e1)** Within CD45.2⁺ donor cells, lineage-negative (Lin⁻) cells were identified, followed by gating on Sca-1 and c-Kit to define the LSK population, and CD48/CD150 expression was used to resolve HSC, MPP, HPC1, and HPC2 subsets. **(e2)** The same lineage, LSK, and CD48/CD150 gating strategy was applied in parallel to CD45.1⁺ competitor-derived cells. This workflow allowed direct quantification of donor versus competitor chimerism across defined HSPC compartments.

